# Photoperiod shapes aluminium tolerance in plants

**DOI:** 10.1101/2021.02.12.430934

**Authors:** João Antonio Siqueira, Thiago Wakin, Willian Batista-Silva, José Cleydson F. Silva, Matheus H. Vicente, Jéssica C. Silva, Wellington R. Clarindo, Agustin Zsögön, Lazaro E. P. Peres, Lieven De Veylder, Alisdair R. Fernie, Adriano Nunes-Nesi, Wagner L. Araújo

**Affiliations:** Departamento de Biologia Vegetal, Universidade Federal de Viçosa; Viçosa, MG, 36570-900, Brazil; National Institute of Science and Technology in Plant-Pest Interactions, Bioagro, Universidade Federal de Viçosa; Viçosa, MG, 36570-900, Brazil; Laboratory of Hormonal Control of Plant Development. Departamento de Ciências Biológicas (LCB), Escola Superior de Agricultura “Luiz de Queiroz”, Universidade de São Paulo; Piracicaba, SP, 13418-900, Brazil; Departamento de Biologia Geral, Universidade Federal de Viçosa; Viçosa, MG, 36570-900, Brazil; Department of Plant Biotechnology and Bioinformatics, Ghent University; Ghent, B-9052, Belgium; VIB Center for Plant Systems Biology; Ghent, B-9052, Belgium; Max-Planck-Institute of Molecular Plant Physiology; Potsdam-Golm, 14476, Germany

**Keywords:** cell-division, DNA-repair, day-length, genetic diversity, photoperiodism

## Abstract

Aluminium is a limiting factor for crop productivity in acidic soils (pH ≤ 5.5). Since acid soil distribution on Earth cannot adequately explain the differential Al tolerance across the plant kingdom, we investigated photoperiod effects on plant Al tolerance. We observed that with increasing distance from the equator, Al tolerance disappears, suggesting a relationship with the photoperiod. Long-day (LD) species are generally more Al-sensitive than short-day (SD) species, whereas genetic conversion of tomato for SD growth boosts Al tolerance. Reduced Al tolerance correlates with DNA-checkpoint activation under LD. DNA-checkpoint-related genes are under positive selection in *Arabidopsis* accessions from regions with shorter days, suggesting photoperiod acts as a selective barrier for Al tolerance. Our findings revealed that diel regulation and genetic diversity affect Al tolerance, suggesting that day-length orchestrates Al tolerance.

**One-Sentence Summary:** Aluminum is a major constraint for crop yield worldwide. We reveal that photoperiod acts as a barrier for Al tolerance in plants.

## Main text

Acidic soils (pH ≤ 5.5) promote the release of aluminium cations, imposing serious constraints on root development in farmland soils around the world. They not only impair nutrient and water uptake (*1*, *2*) but also hinder food production which is required to meet the demands of the growing global population. Notably, soil acidity is also a chronic problem in the temperate zones of eastern North America and throughout Europe (where acidic soils cover up to 80% of the total area). More than 50% of the potentially available arable land in the world is covered by acidic soils (*2*), which results in significant losses in crop productivity. Mechanisms that allow plants to cope with Al stress have previously been elucidated with alterations in organic acid (OA) metabolism (*1*-*3*) and modifications of DNA-repair checkpoints (*4*), which are important in this respect. That said, when analysed individually, neither approach has been demonstrated to be sufficient to overcome Al toxicity. For example, Al-sensitive plants exudate large amounts of OA from root cells to the rhizosphere (*5*), whereas genetic manipulation of the DNA checkpoint machinery does not mediate plant survival in the presence of high Al concentrations (*6*). These findings reinforce the need to identify factors capable of regulating the overall plant response to Al. Furthermore, in the soils of Africa, a large pH range (from alkaline to extremely acidic) is found (Fig. S1) and photoperiod is a dominant factor that controls vegetation phenology and growing season (*7*).

An empirical correlation between the low-pH soils and mean annual day-length amplitude was noted, revealing that the centre of origin of cultivated plants exhibiting lower variations in day-length favour the natural Al tolerance across the plant kingdom (Fig. S1). Likewise, previous studies indicate that most Al-tolerant species are short-day (SD) plants (e.g., *Oryza sativa*, *Stylosanthes guianensis*, and *Vigna unguiculata*), while long-day (LD) plants (e.g., *Hordeum vulgare*, *Pisum sativum*, and *Lens culinaris*) are generally Al-sensitive (Table S1). On analysing the centre of origin of most cultivated plants, we hypothesised that day-length is a pivotal agent modulating Al tolerance across distinct plant species.

## Results

### SD favours Al tolerance regardless of flowering pathway

Since it was observed that photoperiod may play a role in Al tolerance, we hypothesised that an endogenous system also modulates Al-responses. Different leguminous species of plants, some that flower under SD conditions (*Stylosanthes guianensis*, *Vigna unguiculata*, and *Lupinus albans*) and others that flower under LD (*Crotalaria juncea*, *Pisum sativum*, and *Lens culinaris*), were thus selected and cultivated in the presence of Al. It was observed that root elongation in SD species, which are generally Al-tolerant, was insensitive to photoperiod variations, showing similar root elongation rates under both SD and LD conditions, regardless of the presence of Al (Fig. S2). Meanwhile, LD plants, which are usually Al-sensitive, displayed higher root growth and Al sensitivity under LD than under SD conditions, revealing that these plants are indeed more sensitive to Al (Fig. S2). It was further investigated whether fluctuations in day-length altered the mineral-nutrient concentration. Our results revealed that photoperiod influenced Al uptake as revealed by higher Al levels in plants cultivated under SD conditions compared to those cultivated under LD (Fig. S3-4). Intriguingly, even though SD led to a higher Al-tolerance, higher levels of Al were observed in plants growing on SD than under LD, while the levels of other nutrients were not affected by day-length (Fig. S4).

Tomato (*Solanum lycopersicum*) is a day-neutral species, whereas its wild relative *S. pennellii* is an SD species (*8*). The photoperiod-neutrality of *S. lycopersicum* is caused by the loss of function in *SELF-PRUNING 5G* (*SP5G*), which is a flowering repressor that acts on the florigen paralog *SINGLE FLOWER TRUSS* (*SFT*) (*8*). We analysed near-isogenic lines harbouring the *S. pennellii* allele of *SP5G* (*SP5G*^*pen*^) or a loss-of-function mutation in *SFT* (*sft*) in tomato cv. ‘Micro-Tom’ (MT). Based on shoot apical meristem (SAM) analyses, we observed that *SP5G*^*pen*^ modified *S. lycopersicum* into an SD plant, whereas *sft* modified it into a photoperiod-insensitive variety (Fig. S5). Additionally, *SP5G*^*pen*^ plants exhibited a higher Al tolerance, as revealed by similar root growth under both SD and LD conditions and in the presence of Al, a response that was not observed in MT and *sft* plants (Fig. S6-S12). Thus, Al tolerance in tomato seemed to depend on its ability of roots to respond to photoperiod, and not of flowering dependent of day-length.

### Long-term Al tolerance occurs under SD

To further explore the molecular connections between photoperiodic response and Al toxicity, we cultivated *Arabidopsis thaliana* (L.) Heynh. ecotype Columbia 0 wild-type (WT) under SD (8h light/16h dark), neutral days (ND - 12h light/12h dark) and LD (16h light/8h dark), in the presence and absence of Al. Al-induced root growth inhibition was photoperiod-dependent, as neither SD nor ND reduced root growth in the presence of Al. Whereas, under LD, sensitivity to Al-toxicity was enhanced (Fig. S13). It was further investigated whether day-length mitigated reduction in root elongation following either pH change or differential Al concentrations. SD did not ameliorate the reduction in root elongation evoked at lower pH (4.0) compared to the optimal pH (5.7) (Fig. 1A). SD plants were also able to tolerate higher Al levels, showing lower reductions in root elongation than LD plants following increased Al levels (Fig. 1B). Consequently, a multi-level Al response was reported for *A. thaliana*. In addition, following long-term Al exposure, loss-of-function mutants for major DNA-checkpoint regulator genes were generally Al-tolerant, but showed a slower growth recovery after a short-term stress imposed by Al (*6*). In view of this, we further investigated if SD would mitigate plant growth losses resulting from long-term Al exposure. No difference in rosette growth or root elongation was observed in SD plants regardless of Al exposure (Fig. 1C), indicating a likely permanent Al tolerance under SD.

**Fig. 1.**
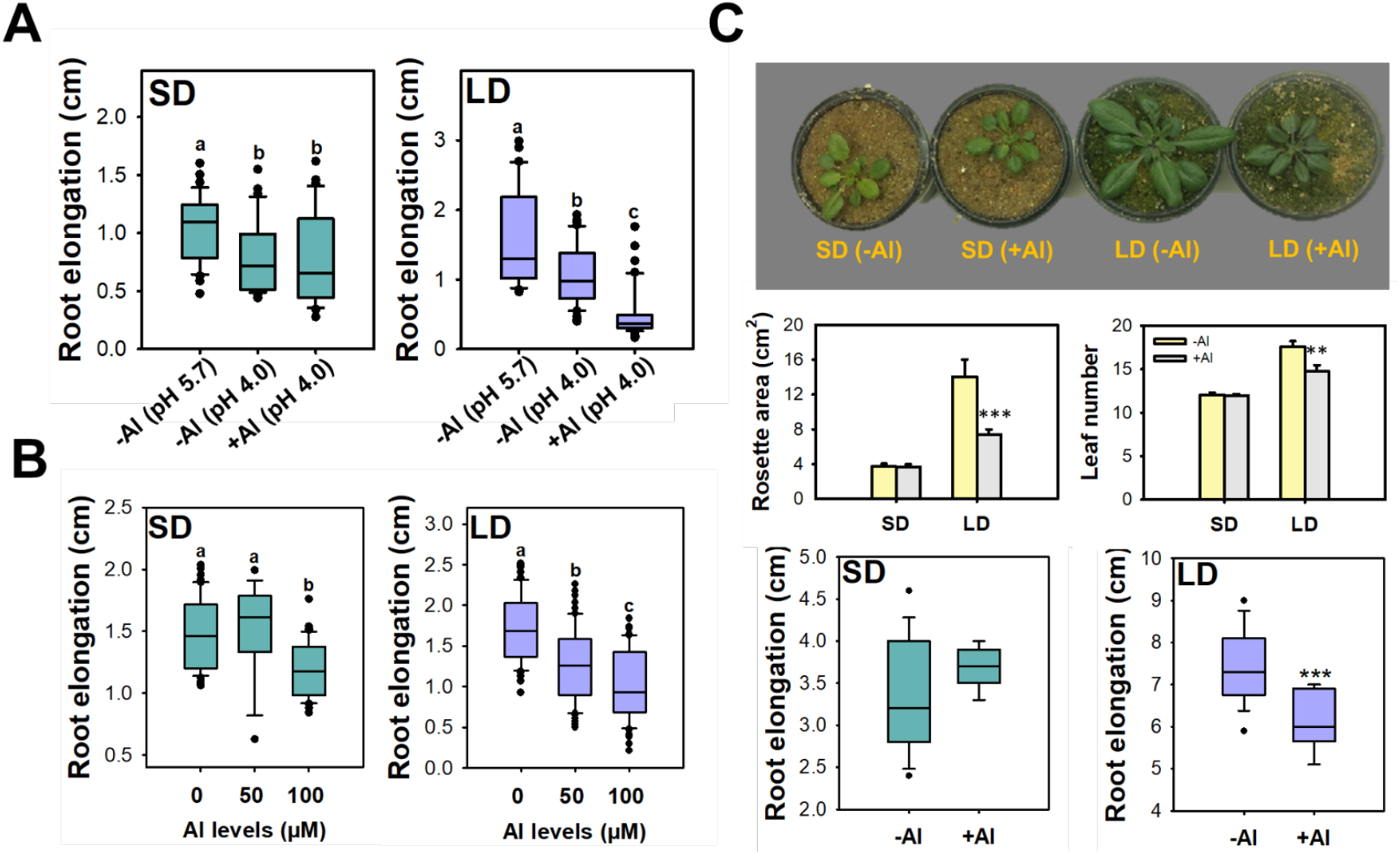
Higher severity of aluminium (Al)-toxicity under long-days (LD) in *Arabidopsis thaliana*. (**A**) Root elongation determined in seedlings growing at pH 5.7 (control) or 4.0 after 10 days, the seedlings were either cultivated with 0 (−Al) or 50 (+Al) μM AlCl_3_ (n = 60 from three independent experiments). (**B**) Root elongation determined in seedlings growing at differential levels of AlCl3 for 10 days (n = 60 from three independent experiments). Statistical groups were determined using a Tukey honest significant difference (HSD) test (*P* <0.05) and are indicated with letters. (**C**) Phenotype for Arabidopsis plants under Al treatment. Representative images of 3-week-old, short-day (SD) or long-day (LD) grown plants cultivated on sand in the absence [–Al (0 μM)] or the presence of Al [+Al (50 μM)]. Measurements from rosette area (cm^2^), leaf number and root elongation for *A. thaliana* plants cultivated on sand (n = 30). Asterisks indicate values that were determined by Student’s *t*-test to be different at *P* < 0.01 (**) or *P* < 0.001 (***) between Al levels.

### Photoperiod specifically mitigates Al toxicity

Cell cycle arrest and DNA damage resulting from Al toxicity are two major cellular alterations affecting plant growth. Al stress culminates in cell cycle blockage and ultimately alters root DNA endoploidy that induces differentiation of root apical meristematic (RAM) cells, a process known as meristem exhaustion (*9*,*10*). Spatiotemporal control of DNA endoploidy was demonstrated across root tissues, indicating an elevated endoploidy in roots coping with low pH (4.6) (*11*). Thus, we isolated RAMs to assess the DNA ploidy levels in response to Al stress, and monitored cell divisions in the root meristems of young seedlings using flow cytometry. Our results revealed the maintenance of potential proliferative capacity (2C – 32C cells) in roots exposed to Al-toxicity under SD, but not under LD conditions. It was also observed that SD suppressed the appearance of polyploid cells (4C, 8C, 16C, and 32C) on the RAM (Fig. 2A), indicating that Al tolerance under SD is likely associated with the downregulation of endoreduplication. Endoreduplication promotes an increase in the DNA ploidy level in several cell types due to changes in cell cycle control. It only occurs in metabolically active and highly specialised cells and allows DNA replication in the absence of mitosis, which increases the levels of DNA endoploidy and regulates root-cell fates (*12*). Our results revealed that under SD, there was a strong reduction in the endoreduplication index in the presence of Al, but it was not so under LD (Fig. 2B). We next turned our attention to identify whether SD promoted reductions in endoreduplication, specifically triggered by Al toxicity, or whether it was a general mechanism of roots under genotoxic conditions. Previous studies have demonstrated that endoreduplication in quiescent-center (QC) cells of the RAM disrupt cell-cycle progression, which might be attributed to the effects of Al toxicity or drugs such as hydroxyurea (HU), methyl methanesulfonate (MMS), and zeocin, leading to a reduced root elongation phenotype (*10*, *13*, *14*). Day-length did not mitigate root elongation limitations triggered by HU, MMS, or zeocin (Fig. 2C), supporting our notion that photoperiod acted specifically in endoreduplication resulting from Al toxicity.

**Fig. 2.**
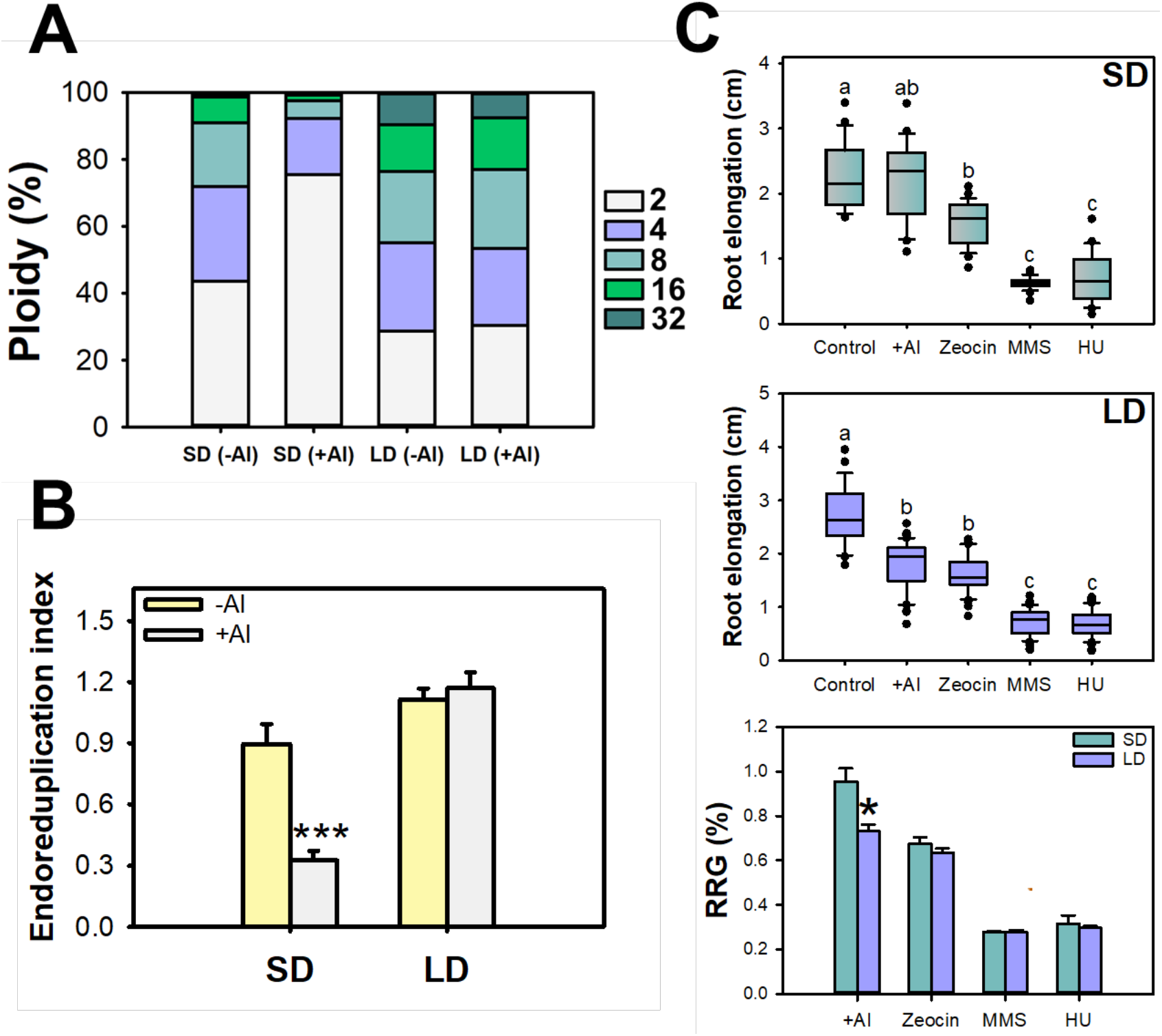
Differential DNA ploidy level and cell cycle regulation modulated by day-length on *Arabidopsis thaliana* seedlings growing under aluminum (Al) stress. DNA ploidy level (%) (**A**) and endoreduplication index (**B**) from root apical meristem (RAM) cells of 10-day-old, short-day (SD)- or long-day (LD) grown seedlings cultivated on – Al (0 μM) or +Al (50 μM). Data represent measurements for five replicates obtained with cells of approximately 30 RAM. (**C**) Seedlings (10-day-old) were submitted to +Al (50 μM), zeocin (5 μM zeocin), methyl methanesulfonate (MMS) (50 ppm) or hydroxyurea (HU) (1 mM), and root elongation (cm) was determined (n = 60 from three independent experiments). RGR (%) indicated root elongation in plants growing under cytotoxic and genotoxic treatments either under SD or LD. RGR (%) was calculated as followed: SD (−Al) root elongation (cm) / SD (treatment) root elongation (cm) (green bars); and LD (−Al) root elongation (cm) / LD (treatment) root elongation (cm) (blue bars). Asterisks indicate values that were determined by Student’s *t*-test to be different at *P* < 0.05 (*) or *P* < 0.001 (***) between Al levels. Statistical groups were determined using a Tukey honest significant difference (HSD) test (*P* value <0.05) and are indicated with different letters.

### LD are required to arrest cell divisions under Al stress

Endoreduplication is an alternative pathway that avoids cell cycle arrest or cell death due to DNA damage (*15*). Thus, we investigated how SD promotes reductions in endoreduplication during Al toxicity. In response to Al, endoreduplication is mainly modulated by the transcription factor SUPPRESSOR-OF-GAMMA-RESPONSE1 (SOG1), which is activated by the kinases ATAXIA-TELANGIECTASIA-MUTATED (ATM) and ATAXIA-TELANGIECTASIA-AND-RAD3-RELATED (ATR), enhancing the expression of genes associated with cell cycle stoppage and DNA repair (*10*). *A. thaliana* loss-of-function mutants for *SOG1* and *ATR* displayed higher Al tolerance due to the inhibition of early endoreduplication onset (*10*). *In silico* analyses further revealed that most of the genes involved in DNA repair and endoreduplication regulation exhibited a significant correlation with latitude/day-length in 32 *A. thaliana* accessions (Fig. S14). In agreement with our *in silico* analyses (Fig. S15), qRT-PCR analysis of the major DNA-repair regulators (*SOG1*, *ATM*, and *ATR*) revealed a more coordinated gene expression under SD than under LD conditions (Fig. 3). Briefly, *SOG1*, *ATM*, and *ATM* expression increased in a light-dependent manner only during LD, reaching maximum expression peaks at points near dusk (Fig. 3). Based on the contribution of both SOG1 and ATR checkpoint regulators on the Al-toxicity response, it was hypothesised that their diel regulation might have contributed to the observed photoperiodic Al-tolerance. We confirmed this by growing *A. thaliana* mutants defective in DNA checkpoint control, *sog1-1*, *atm-1*, and *atr-2*. As expected, *sog1-1*, *atm-1*, and *atr-2* showed an invariant Al tolerance following day-length variations, with similar root growth responses following SD and LD (Fig. S16). Collectively, SD seemed to support mitosis maintenance by repressing excessive DNA replication that would culminate in higher endoreduplication.

**Fig. 3.**
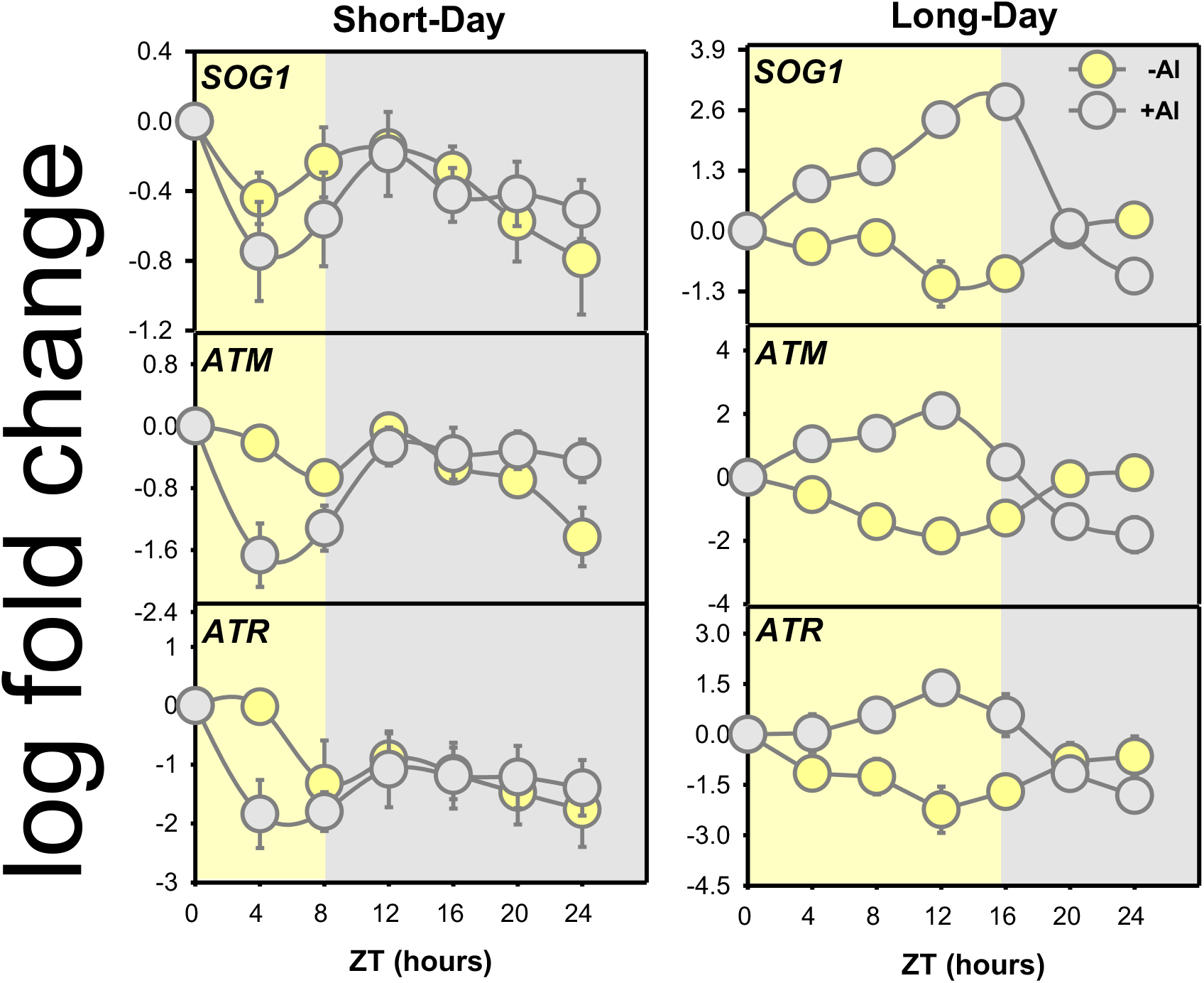
Expression profile of regulators of DNA checkpoints is variable in response to photoperiod and Al. Diurnal oscillations of transcript levels from *SOG1* (*SUPPRESSOR OF GAMMA RESPONSE1*), *ATM* (*ATAXIA TELANGIECTASIA MUTATED)* and *ATR* (*ATAXIA TELANGIECTASIA AND RAD3 RELATED*) were determined in roots of 3-week-old plants of *Arabidopsis thaliana* ecotype Columbia (Col-0) cultivated at −Al (0 μM) or +Al (50 μM) under short-days (SD) and long-days (LD). Light and dark rectangles denote day and night periods in SD (8 h light/16 h dark) and LD (16h light/8h dark), respectively. Data represent the average expression of three biological replicates ± SE.

### Circadian clock and genetic diversity impose constraints for Al-tolerance

Day-length is a more predictable, unperturbed, and non-oscillating factor differing from temperature, air humidity, and rainfall. Thus, *A. thaliana* accessions from high-latitude environments usually experience long photoperiods around summer solstice, exhibiting longer circadian periods (*16*). Casein kinase 2 (CK2) phosphorylates both circadian clock proteins related to the light cycle (*17*) and SOG1 protein on amino acid T423 (*18*). Correspondingly, the mutation of T423 into its phosphomutant A (T423A) mimics the Al-resistant phenotype of *A. thaliana sog1-101* mutant plants (*18*). Interestingly, *sog1-101* and *T423A-27* knockout mutants displayed root elongation insensitive to day-length as well as higher Al-tolerance than WT plants, regardless of the photoperiod (Fig. S17). Therefore, we postulate that genetic diversity could occur in genes related to DNA and cell cycle in *A. thaliana*, which would be selected in response to day-length and light components of the circadian clock machinery. In keeping with this assumption, we analysed two independent studies (*19*, *20*) to verify whether latitudinal/day-length variation affected root elongation in *A. thaliana* accessions. Interestingly, we observed a positive and significant correlation between latitude/day-length and root length, revealing that accessions from higher latitudes displayed greater root length (Fig. S18). Regarding root development, only with light incidence on shoots, cell death was triggered around the RAM cells of *A. thaliana* seedlings (*21*). Furthermore, to identify the existence of genetic diversity for genes investigated here as members of the interface between Al-tolerance and day-length, we selected 287 *A. thaliana* accessions with the centre of origin regions widely varying in photoperiod. By grouping approaches, we found three distinct groups (G1, G2, and G3) based on day-length in the region origin, in which G1 exhibited more accessions with photoperiod around 12-14h while G3 had the majority of accessions from regions with day-length of around 20h (Fig. 4A-B). Assessing the relatedness of *A. thaliana* accessions using STRUCTURE, we were also able to discriminate individuals from three populations (Fig. 4C), revealing an essential absence of a mixture of accessions due to photoperiod-barriers. Therefore, the higher *fitness* of *A. thaliana* accessions from low-latitudes under drought conditions (*22*) and more coordinated oscillatory behaviour in drosophilids (*23*) might be due to the lower photoperiod variability across the year. Likewise, the mapping of polymorphisms in genes involved in DNA repair, cell cycle, and endoreduplication revealed the existence of a characteristic genetic diversity for genes involved in Al tolerance in *A. thaliana*, which varied with day-length (Fig. S19). Furthermore, we found more non-synonymous and synonymous mutations in *ATM*, *ATR*, and *SOG1* genes in accessions derived from longer day regions (G3), the unique group in which these mutations were correlated with day-length (Fig. 4D-E and S20). In agreement with these findings, the global distribution of population genetic diversity across the animal and plant kingdoms reveal that only eudicots exhibit a significant correlation between population genetic diversity and latitude Notably, genetic diversity increases with distance to the equator within eudicots (*24*). Strikingly, accessions from lower day regions (G1 and G2) were under positive selection (*dN*/*dS* ≥ 1), meaning that the selective patterns are directly related to gene expression (Fig. 4F, S14, and S20). Therefore, photoperiod seems to have been imposing selection patterns in genes that are known to improve Al tolerance. Collectively, our results suggested that photoperiod is likely a factor that mediates plant Al tolerance.

**Fig. 4.**
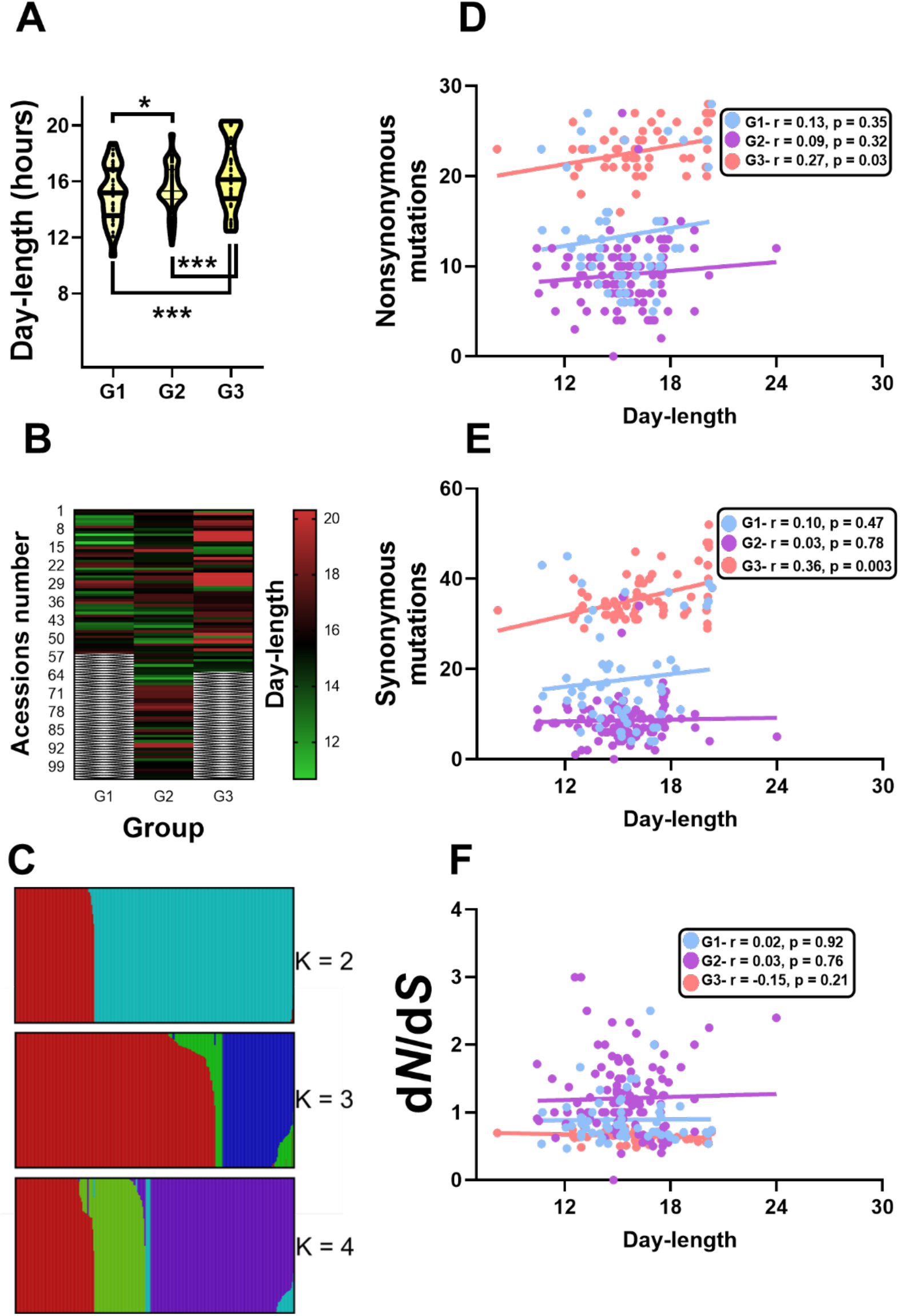
Photoperiods discriminate *Arabidopsis thaliana* accessions according with genetic variants of genes related to cell cycle and DNA checkpoints. 287 *A. thaliana* accessions from centre of origin varying in day-length from a duration of 8:11 to 23:53 were selected based on previous studies. (**A**) Violin-plots representing group analyses were clustered by the Orange Canvas software and a discriminant analysis of principal components, in which three major accession groups G1 (n = 59), G2 (n = 112) and G3 (n = 63) based on distinct day duration were formed. Remarkably, some accessions were not grouped into these three groups. Asterisks indicate values that were determined by Student’s *t*-test to be different at *P* < 0.05 (*) or *P* < 0.001 (***) between groups. (**B**) Heat-map highlighting the differential composition of day-length on groups. (**C**) Population structure of *A. thaliana* accessions in which each vertical bar represents an individual accession with single nucleotide polymorphism (SNP) on genes related to cell cycle and DNA checkpoints. K clusters indicate colours for fractional memberships. Further details concerning genes and accessions used in the analyses can be found in the supplementary material. (**D-F**) Accumulation of non-synonymous and synonymous mutations as well as the ratio of non-synonymous to synonymous fractions (*dN*/*dS*) and its correlations with day-length. By using the POLYMORPH 1001 (https://tools.1001genomes.org/polymorph/), polymorphism was detected on genes *ATAXIA TELANGIECTASIA MUTATED* (*ATM*), *ATAXIA TELANGIECTASIA AND RAD3 RELATED* (*ATR*), and *SUPPRESSOR OF GAMMA RESPONSE1* (*SOG1*) among groups G1, G2, and G3.

## Discussion

Our results suggested a crucial yet unreported significance of photoperiod and endogenous cues, which appear to modulate Al tolerance across the plant kingdom. Our data further indicated that SD is most likely associated with higher Al-tolerance. Intriguingly, SD plants could uptake higher Al levels than LD plants. Reprogramming of the mitochondrial OA metabolism may mediate Al tolerance in both microorganisms and plants (*1*-*3*). Higher Al content in the roots of plants growing under SD indicates an internal immobilisation of Al in the apoplast. Further work is clearly required to assess the significance of biochemical changes in photosynthesis and respiration, which are the major processes that produce and consume photoassimilates. Our genetic diversity mapping also suggested that (*i*) the lower competition among plant species at higher latitudes may enable the occurrence of species adapted to stressful conditions, and (*ii*) with increasing distance from the equator, stressful environmental conditions may alter patterns of energy allocation from vegetative growth to reproduction, increasing the genetic diversity in populations from these regions (*24*,*25*). Day-length specifically influences cell cycle arrest and endoreduplication related to Al, modulating gene expression in a diel manner. Since a genetic variability for these genes was observed according to photoperiod, it seems reasonable to question whether day-length regulates only Al responses or it could coordinate tolerance to other abiotic stresses as well. In addition, our results offer novel perspectives for understanding the underlying mechanisms of Al-tolerance in land plants. For example, under field conditions where plants are recurrently exposed to fluctuating conditions, Al-tolerance mechanisms involving different levels of complexity can play a pivotal role in helping plants to successfully cope with high levels of Al. Additional studies, which employ more sophisticated and combined analyses, will likely be of fundamental importance in providing a holistic understanding of the underlying mechanisms that affect the evolution of Al-tolerance. Therefore, our findings suggest a wide yet unrecognised significance of day-length in orchestrating Al tolerance, which might enable us to develop the next generation of productive crops under higher Al conditions.

## Acknowledgments

We thank Dimas Mendes Ribeiro and Cleberson Ribeiro for helpful comments and sound advice on an earlier draft of this manuscript.

## Author contributions

J.A.S., and W.L.A. designed the research; J.A.S. performed most of the research with the support of T.W. and W.B-S.; J.C., and W.C. performed cytometry flow analyses; J.A.S., and J.C.F.S. realized bioinformatics analyses; W.B-S., J.C., W.C., L.D.V., A.R.F., and A.N-N. contributed new reagents/analytic tools; J.A.S., L.D.V., A.N-N., and W.L.A. analysed the data; and J.A.S., and W.L.A. wrote the article with input from all the others.

## Data and materials availability

All data are available in the main text or the supplementary materials.

## Supplementary Materials

Materials and Methods

Figs. S1 to S20

Tables S1 to S3

References

Data S1 to S3

