## Supplementary for "Photoperiod shapes aluminium tolerance in plants"

**This PDF file includes:**

Materials and Methods
Figs. S1 to S20
Tables S1 to S3
References

**Other Supplementary Materials for this manuscript include the following:**

Data S1 to S8 [Data S1_Num_PCA_Retained, Data S2_PCA, Data S3_Pop_Orange_Arabidopsis, Data S4_scatter, Data S5_barplot, Data S6_compoplot, Data S7_Pop_Orange_Arabidopsis, and S8 Arabidopsis thaliana accessions details]

**Materials and Methods**

Plant material and growth conditions

Different plant species were used in this study. Briefly, we used: (i) Arabidopsis wild-type (WT) and mutant plants of DNA repair and cell cycle related genes, of the Columbia-0 ecotype (Col-0); (ii) tomato (*Solanum lycopersicum* 'MT') and loss of function mutants in the *SELF-PRUNING 5G* (*SP5G*) and *SINGLE FLOWER TRUSS* (*SFT*) as well as near-isogenic lines harbouring the *S. pennellii* allele of *SP5G* (*SP5G*$^{pen}$); and (iii) different leguminous species, namely *Stylosanthes guianensis*, *Vigna unguiculata*, and *Lupinus albans*, *Crotalaria juncea*, *Pisum sativum,* and *Lens culinaris*. Further details are provided in Table S2. Briefly, seeds were surface sterilized by agitation in 30% (v/v) commercial bleach (2.7% [w/v] sodium hypochlorite) for 15 min, followed by three rinses with sterile distilled water and kept in darkness to synchronise germination for 4 days. Seedlings of all plant species, except *Arabidopsis thaliana*, were cultivated in hydroponics solution by using Hoagland medium (Hoagland and Arnon, 1950) at pH 4.0, with modifications. Plants were grown in a full-strength solution (pH 4.0) with 100 µM $AlCl_3$ (+Al) and 0 µM (-Al) for 5 days, with pH adjusted daily to 4.0. Plants were cultivated in a temperature-controlled chamber (20 ± 1 °C for Arabidopsis, and 25 ± 1°C for other species) under 250 µmol photons $m^{-2}$ $s^{-1}$, 60% relative humidity, and either under short-days (SD: 8-h light/16-h dark) or long-days (LD; 16-h light/8-h dark). Root systems were photographed, and root elongation was measured using the ImageJ software (https://imagej.nih.gov/ij/); root systems were harvested at 5 days after sowing (DAS) and further used for nutritional analyses. To investigate the long-term impacts of Al exposure, we transferred 10-day-old *in vitro-grown A. thaliana* seedlings to pots containing washed sand. These plants were cultivated for an additional two weeks and received 7 mL of Murashige and Skoog (MS medium) solution daily, with Al or +Al (50 µM $AlCl_3$) at pH 4.0.

For *in vitro* assays, seeds from *A. thaliana* were surface-sterilized and imbibed for 4 days at 4°C in the dark on 0.8% (w/v) agar plates containing half-strength MS medium (Sigma-Aldrich; pH 4.0), with different $AlCl_3$ concentrations (0, 50, and 100 μM). Next, seedlings were cultivated for 10 days in a growth chamber (POL-EKO APARATURA® Climatic Chamber KK 1200) under SD and LD at 22 ± 1°C, 60% relative humidity, and 150 μmol photons $m^{-2}$ $s^{-1}$. In a second experiment, the steps described above were followed and in addition to the treatments mentioned, we used MS medium at pH 4.0 supplemented with zeocin (5 μM zeocin), methyl methanesulfonate (MMS) (50 ppm), or hydroxyurea (HU) (1 mM). Root elongation was determined *in vitro* in a similar manner to the hydroponics experiments.

Flow cytometry

Roots from *A. thaliana* seedlings were collected, and the root apical meristems (RAM) were excised. From ~30 RAM for each repetition, nuclei were isolated using grinding movements with a pestle in 0.2 mL OTTO-I lysis buffer (Otto 1990) supplemented with 2.0 mM dithiothreitol (Sigma®), and then made up to 0.8 mL with the same buffer. The nuclei suspensions were then filtered through a 20 μm nylon mesh (Partec®) and centrifuged at 100*g* × g for 5 min. The obtained pellet was incubated for 10 min in 0.1 mL of OTTO-I lysis buffer and stained with 0.5 mL of OTTO-I:OTTO-II (1:2) solution, supplemented with 75 μM propidium iodide and 2.0 mM dithiothreitol (Sigma®). The nuclei suspensions were incubated for 30 min in the dark at room temperature. Each suspension was analysed using a BD Accuri C6 flow cytometer (Accuri cytometers, Belgium) equipped with a 488 nm laser source. FL2 (585/640) and FL3 (670 LP) filters were used to detect propidium iodide fluorescence. The BD Csampler software (Accuri Cytometers, Belgium) was used for histogram analyses to determine the DNA ploidy level of each

$G_0/G_1$ peak. For this, we considered only the histograms with $G_0/G_1$ peaks exhibiting a coefficient of variation below 5% for the $G_0/G_1$ peak and at least 5,000 nuclei were counted for each nuclei suspension.

Nutrient analyses

To quantify the levels of Al, phosphorus, potassium, calcium, magnesium, the dry root tips (~0.5 cm) were subjected to a nitric-perchloric digestion (65% and 70%) (Miyazama et al. 1999). The samples were analysed using inductively coupled plasma optical emission spectroscopy (ICP-OES, Perkin-Elmer Optima 3000XL, Maryland, USA).

Expression analysis by qRT-PCR

Total RNA was extracted from root samples harvested (immediate snap-freezing in liquid nitrogen) at time points described in the specific figure captions. The RNA was isolated using the TRIzol reagent (Ambion, Life Technology), according to the manufacturer's recommendations. It was then treated with DNase I (RQ1 RNase free DNase I; Promega). RNA integrity was analysed on 1% (w/v) agarose gels by measuring the RNA concentration with a Nanodrop spectrophotometer. Real-time PCR was performed with cDNA using a sequence detection system (Applied Biosystems Applera) using the Power SYBR Green PCR Master Mix according to Piques et al. (2009). The calculations for transcript abundance were performed with standard curves of each selected gene and normalised using the constitutively expressed gene ACTIN (AT2G37620). Data analyses were performed as described previously (Caldana et al., 2007). The primer sequences used are shown in Table S3. Melting curves were checked for unspecific amplification and primer dimerization.

Genetic structure analysis

We selected 287 *A. thaliana* accessions from the 1001 Genomes Consortium database (https://1001genomes.org/) (Alonso-Blanco et al., 2016) based on variable day-length from 8:11 to 23:53 of duration, calculated by http://www.solartopo.com/daylength.htm. Single Polymorphism Nucleotide (SNPs) were identified in genes related to cell cycle and DNA repair by using the POLYMORPH 1001 (https://tools.1001genomes.org/polymorph/). The genetic structure population was analysed using fast structure software (Raj et al., 2014). Unsupervised machine learning approaches were used to analyse the accession groups. The accession data were hierarchically clustered with the Orange Canvas software (http://orange.biolab.si/) using Pearson's correlation distance as the distance measure and the average linkage clustering option. The discriminant analysis of principal components (DAPC) was performed using the 'adegenet' package 1.4-1 (Jombart et al., 2010) in R studio V 2.3.2 (R Development Core Team, 2011) to confirm cluster numbers and to describe the global diversity for overlooking differences between groups.

Statistical Analyses

All experiments were designed in a completely randomised distribution with a minimum of three biological replicates of each treatment. Additionally, the experiments were repeated at least three times (even in different growth facilities) with similar phenotypes observed each time. Data were statistically tested for normality and subsequently examined using ANOVA ($P < 0.05$).

Differences in the means ($P < 0.05$) displayed in figures and tables were examined by Student's *t-test*. All statistical analyses were performed using R statistical software (www.r-project.org).

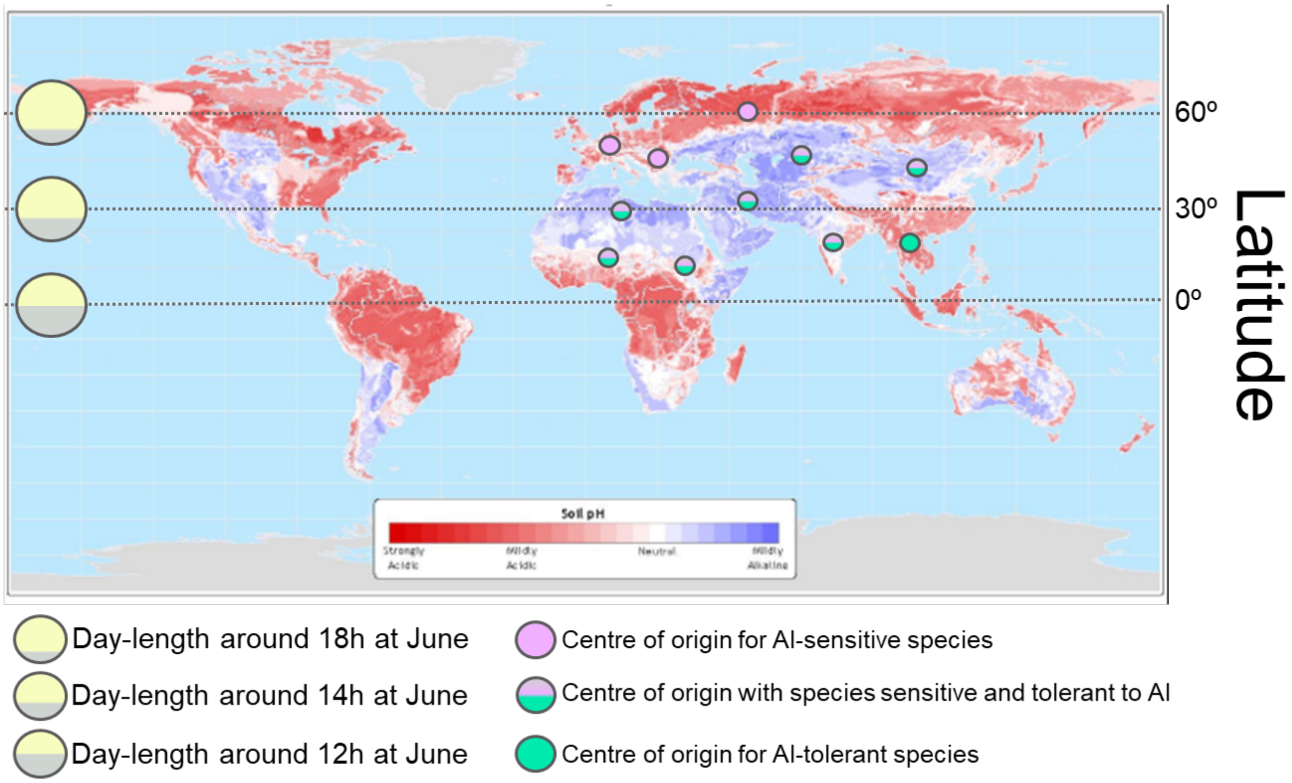


**Supplementary Figure S1. Hypothetical scheme highlighting possible interfaces between photoperiod and aluminium (Al) tolerance.** Apparently, the soil pH at the centre of origin of most plant species is not capable to adequately explain the differences in Al tolerance across the plant kingdom. By looking at the centre of origin of the major cultivated plants, Al tolerance is seemingly lost in species that originated further way from the Equator line. Therefore, photoperiod could explain, at least partially, the differential Al tolerance across the plant kingdom, since many origin centres harbouring Al tolerant plants are situated on regions with alkaline soils, which suggests the pivotal importance of photoperiod to select Al tolerance. Interestingly, centre of origin with higher latitudes, where day-length at summer solstice (June) reach 18 hours or more, are characterized predominantly by the presence of Al sensitive species. Taken together, we hypothesized that photoperiod can modulates plant responses to Al. The map is an adaptation from the Atlas of the Biosphere (Center for Sustainability and the Global Environment, University of Wisconsin, Madison) (https://nelson.wisc.edu/sage/data-and-models/atlas/maps.php).

**Supplementary Table S1. Relationship between aluminium (Al) tolerance levels and photoperiodism.** We observed, across the plant kingdom, plants exhibiting the growing-season at origin centre with short-days (green), neutral-days (orange), and long-days (red). Long-day plants apparently exhibit a tendency to be more sensitive to Al, while short-day plants have propensity to display lower yield alterations under Al stress (higher tolerance). This table was constructed following Meda and Furlani (2005) and Garcia-Oliveira *et al.* (2016).

| Al tolerance degree | Crop species* |
|---|---|
| *Highly sensitive* | **Alfalfa**, **barley**, **carrot**, **durum wheat**, **lettuce** and **pea** |
| *Sensitive* | **Oat** and **Wheat** |
| *Moderately sensitive* | **Cabbage**, **maize** and **sorghum** |
| *Moderately tolerant* | **Rice** and **rye** |
| *Tolerant* | **Soybean** and **pigeon pea** |
| *Highly tolerant* | **Tea**, **buckwheat**, **brachiaria**, **Vigna unguiculata**, and **Stylosanthes** |

Crop species and their scientific names are as follow: Alfalfa (*Medicago sativa*), barley (*Hordeum vulgare*), *Brachiaria spp.*, buckwheat (*Fagopyrum esculentum*), cabbage (*Brassica oleracea* var. capitata), carrot (*Daucus carota* subsp. sativus), durum wheat (*Triticum durum*), lettuce, (*Lactuca sativa*) maize (*Zea mays*), oat (*Avena sativa*), pea (*Pisum sativum*), pigeon pea (*Cajanus cajan*), rice (*Oryza sativa*), rye (*Secale cereale*), sorghum (*Sorghum bicolor*), soybean (*Glycine max*), *Stylosanthes spp.*, Vigna unguiculata and wheat (*Triticum aestivum*).

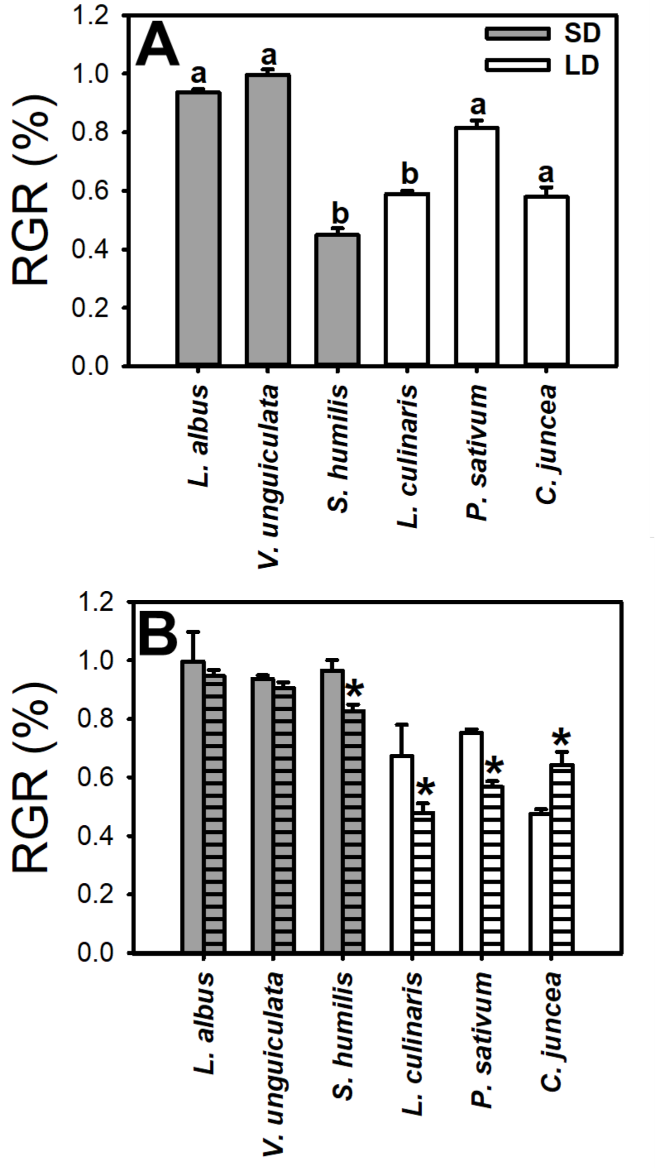


**Supplementary Figure S2. Root elongation and aluminium (Al) tolerance are affected by photoperiod.** (**A**) Relative growth rate (RGR - %) was determined by the root elongation ratio measured in species from short-days (SD, grey) and long-days (LD, white) plants growing at SD and LD conditions under control conditions (absence of Al). Root elongation was determined in 5-day-old seedlings. RGR (%) was calculated as followed: SD root elongation (cm) / LD root elongation (cm) (*n* = 16 plants for each condition). Statistical groups were determined using a Tukey honest significant difference (HSD) test ($P < 0.05$) and are indicated with different letters. (**B**) RGR (%) indicated root elongation measured in plants growing in the absence of $Al^{3+}$ or in the presence of 100 µM $Al^{3+}$ under either SD (clear bar) or LD (hatches bar) conditions. RGR% was calculated as follow: SD (-Al) root elongation (cm) / SD (+Al) root elongation (cm) (clear bars); and LD (-Al) root elongation (cm) / LD (+Al) root elongation (cm) (hatches bar).

An asterisk (*) indicate values that were determined by the two-sided Student's *t*-test to be different ($P < 0.05$) between SD and LD. SD species: *Lupinus albus*, *Vigna unguiculata* and *Stylosanthes humilis*; LD species: *Lens culinaris*, *Pisum sativum* and *Crotalaria juncea*.

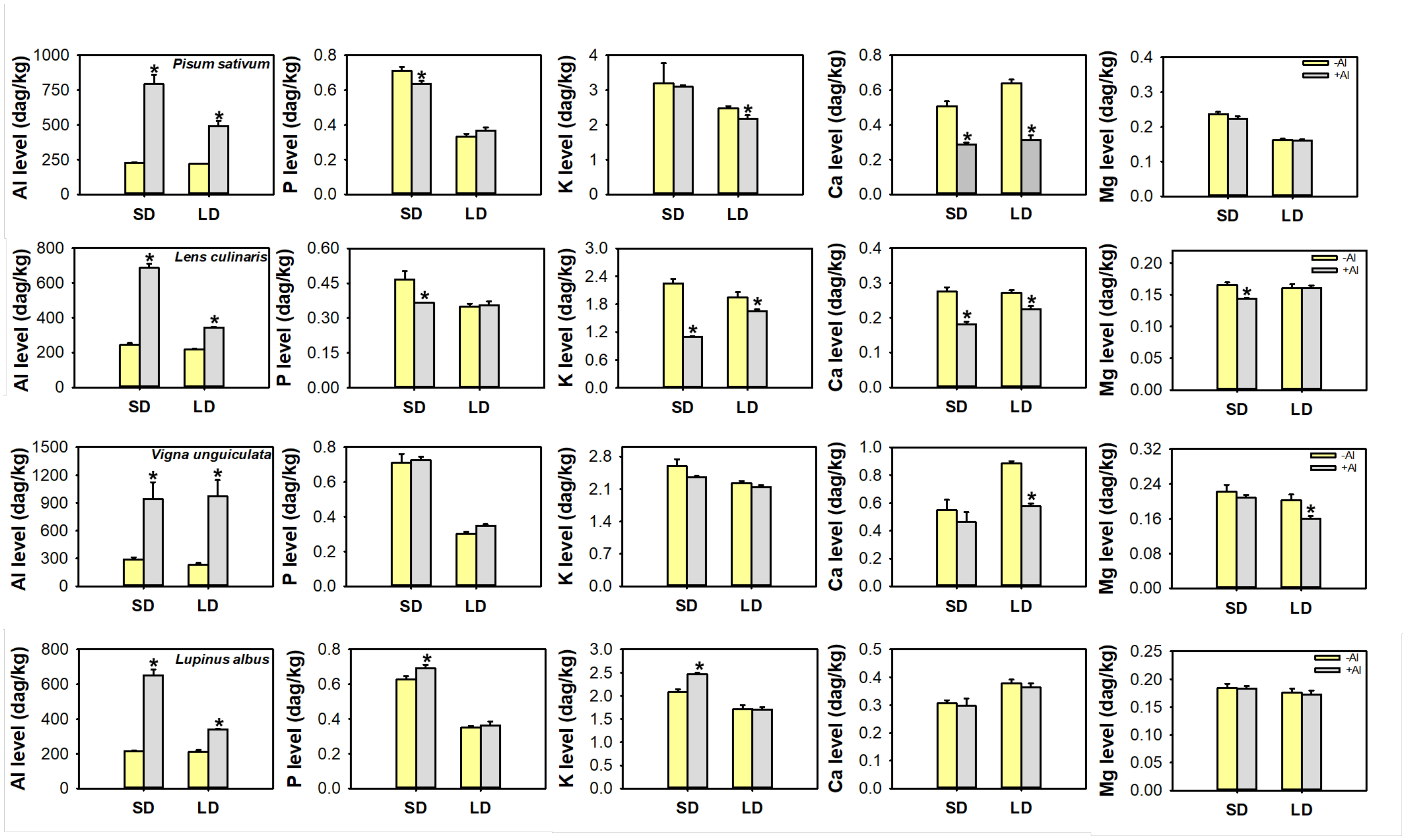


**Supplementary Figure S3. Differential nutritional content in response to changes in photoperiod under Aluminium (Al) stress.** The levels of Al, Phosphorus (P), Potassium (K), Calcium (Ca) and Magnesium (Mg) were determined on root samples (n = 4 samples for each condition) of plants growing at short-days (SD) or long-days (LD). Data represent nutrient levels determined in plants growing in the absence (-Al, yellow) or the presence 100 μM Al (+Al, grey) after five days on hydroponics culture. An asterisk (*) indicate values that were determined by the two-sided Student's *t*-test to be different ($P < 0.05$) between -Al and +Al.

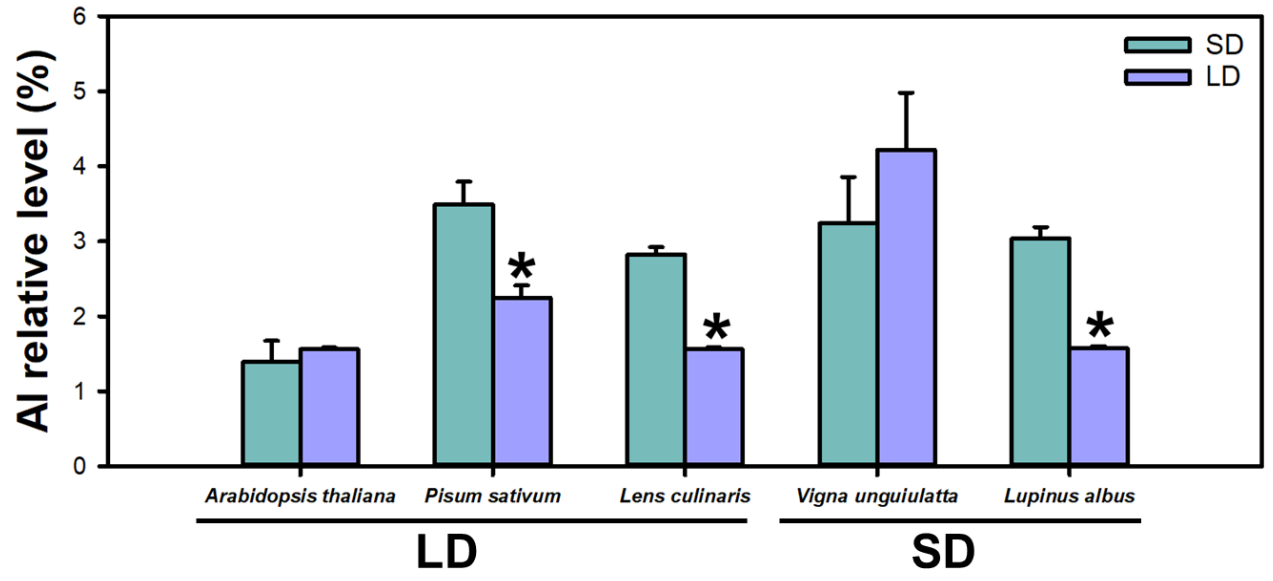


**Supplementary Figure S4. Aluminium (Al) relative levels (%) is differentially affected by Short-days (SD) and long-days (LD).** The levels of Al were determined on root samples (n = 4 samples for each condition) of plants growing at short-days (SD) or long-days (LD) and are additionally shown in Supplemental Figure S3. The relative levels were determined as follow: SD (-Al) Al levels / SD (+Al) Al levels (green bars); and LD (-Al) Al levels / LD (+Al) Al levels (blue bars). The plants were cultivated the absence of $Al^{3+}$ (-Al) or in the presence of 100 µM $Al^{3+}$ (+Al) under SD or LD by five days on hydroponics culture. An asterisk (*) indicate values that were determined by the two-sided Student's *t*-test to be different ($P$ <0.05) between SD and LD.

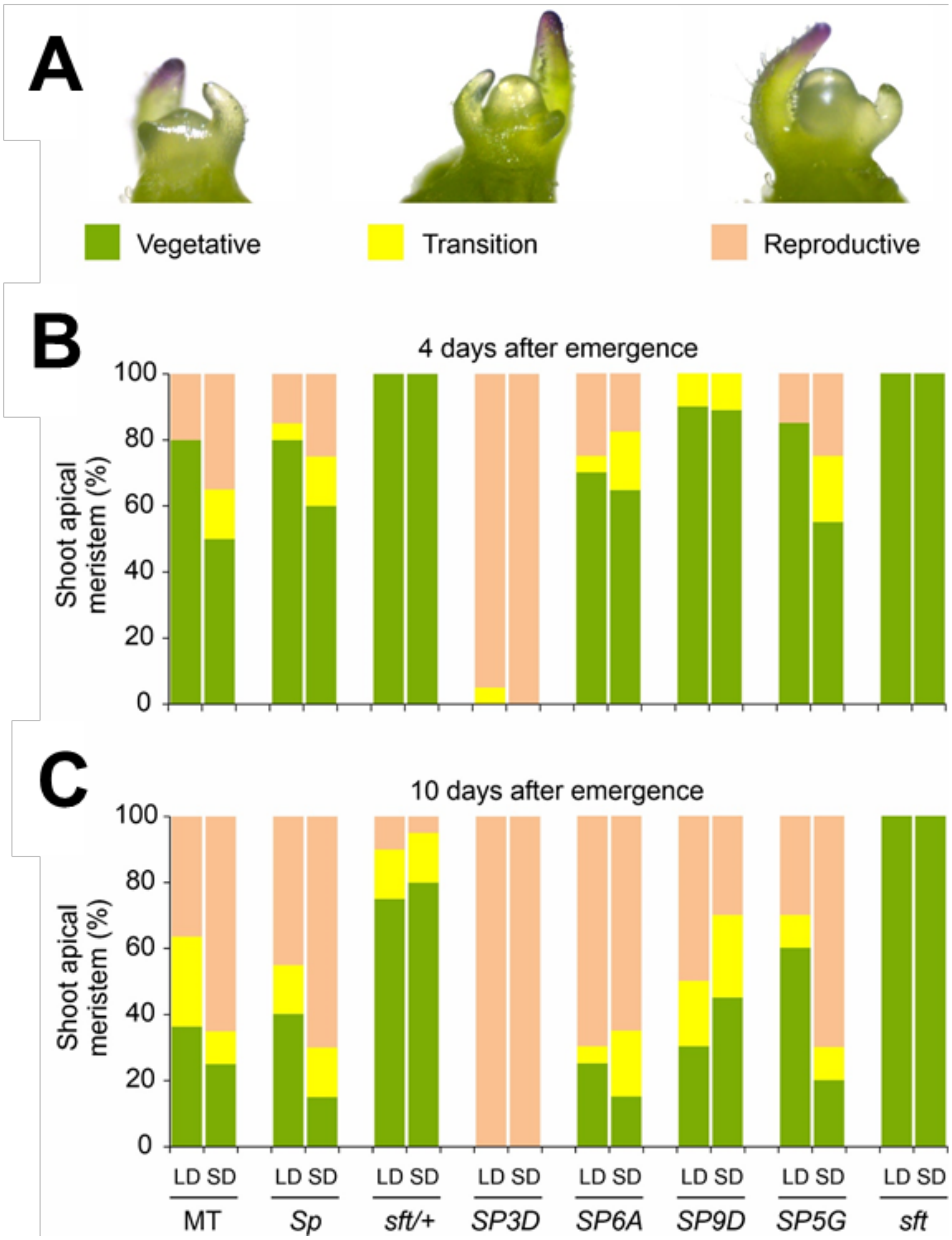


**Supplementary Figure S5. Day-length regulates differently genes of the *SELF-PRUNING* family to orientate shoot apical meristem (SAM) transition**. **A**, Representation of meristem maturation stages of SAM, vegetative (left), transition (center) and reproductive (right). **B-C**, SAM maturation at 4 (B) and 10 (C) days after germination on short (SD) and long (LD) days (n= 17-20 seedlings). Abbreviations: Micro-Tom (MT), *Self-pruning* (*Sp*), *Single flower truss* (*sft*/+ and *sft*), *Solanum pennellii* alleles *SELF-PRUNING 3D*, *6A*, *9D*, and *5G* (*SP3D*, *SP6A*, *SP9D*, and SP5G).

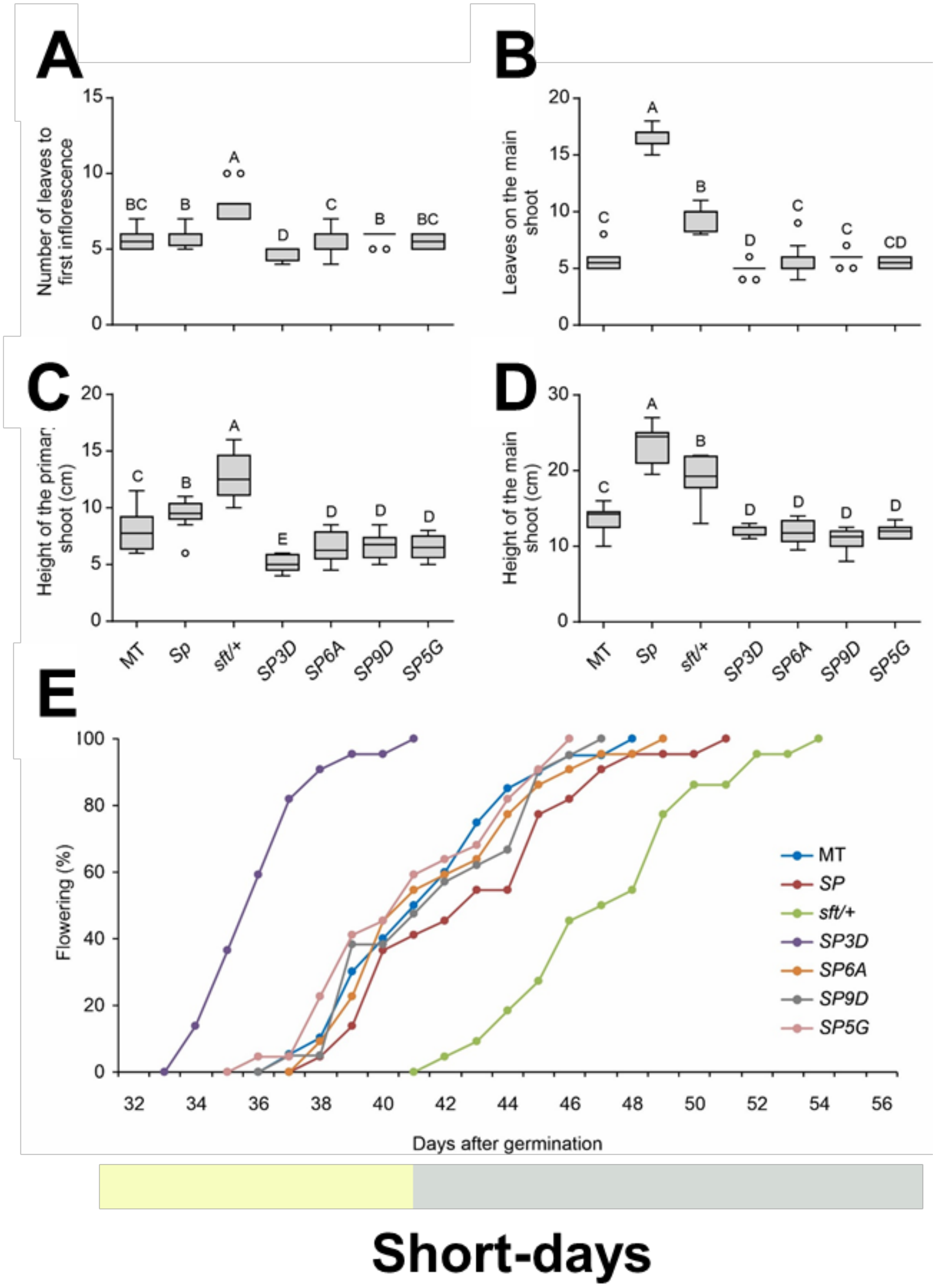


**Supplementary Figure S6. Phenotype of tomato genotypes carrying distinct alleles from the *SELF-PRUNING* gene family under short-days. A**, leaf number until the first inflorescence. **B**, leaf number on the main shoot. **C**, height of the primary shoot, or until the first inflorescence insertion. **D**, height of the main shoot. **E**, flowering time in days after germination. The values represent 20-22 plants per genotype, and statistical groups were determined using a Tukey honest significant difference (HSD) test ($P < 0.05$) and are indicated with different letters.

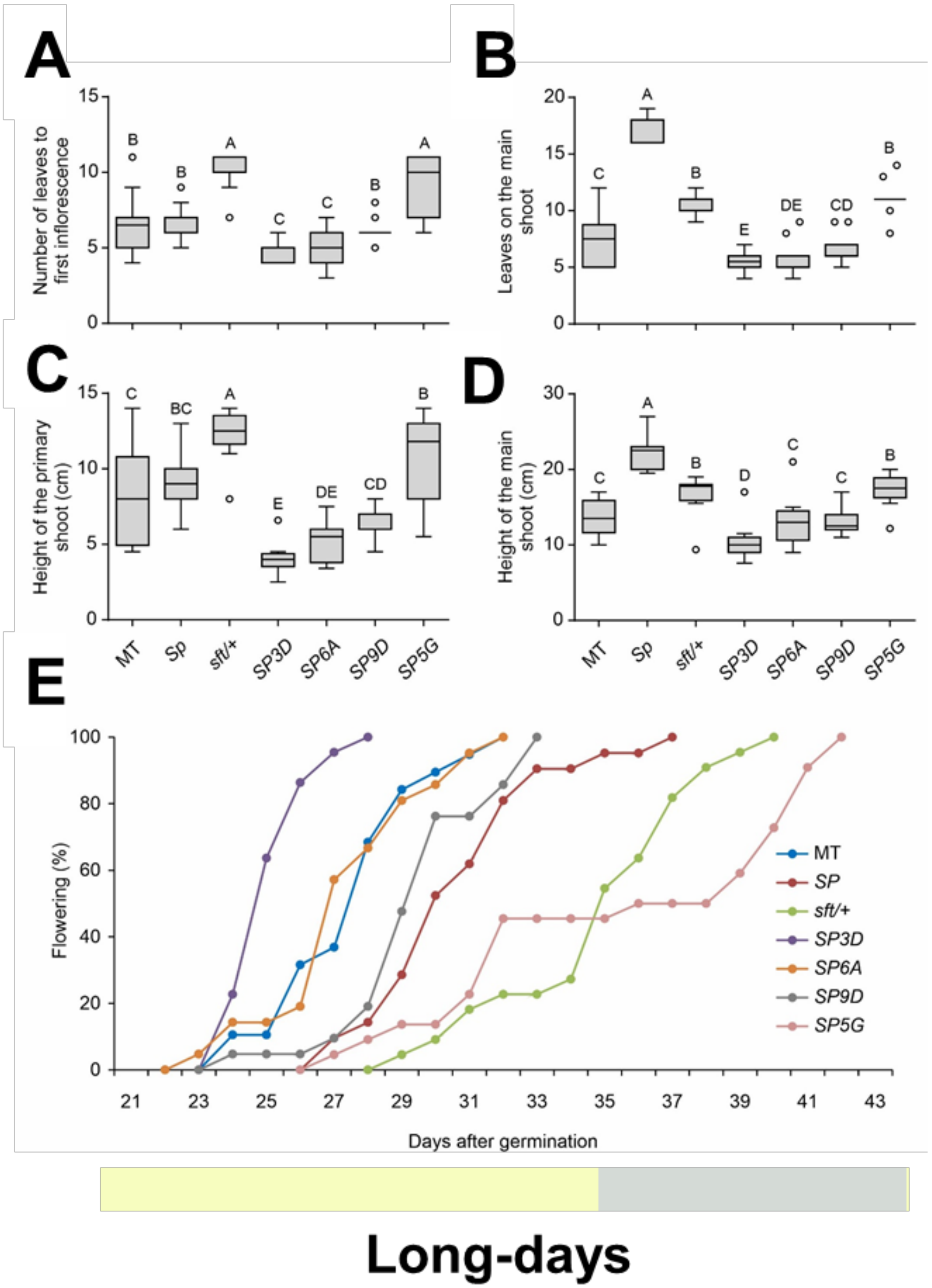


**Supplementary Figure S7. Phenotype of tomato genotypes carrying distinct alleles from the SELF-PRUNING gene family under long-days. A**, leaf number until the first inflorescence. **B**, leaf number on the main shoot. **C**, height of the primary shoot, or until the first inflorescence insertion. **D**, height of the main shoot. **E**, flowering time in days after germination. The values represent 20-22 plants per genotype, and statistical groups were determined using a Tukey honest significant difference (HSD) test ($P$ <0.05) and are indicated with different letters.

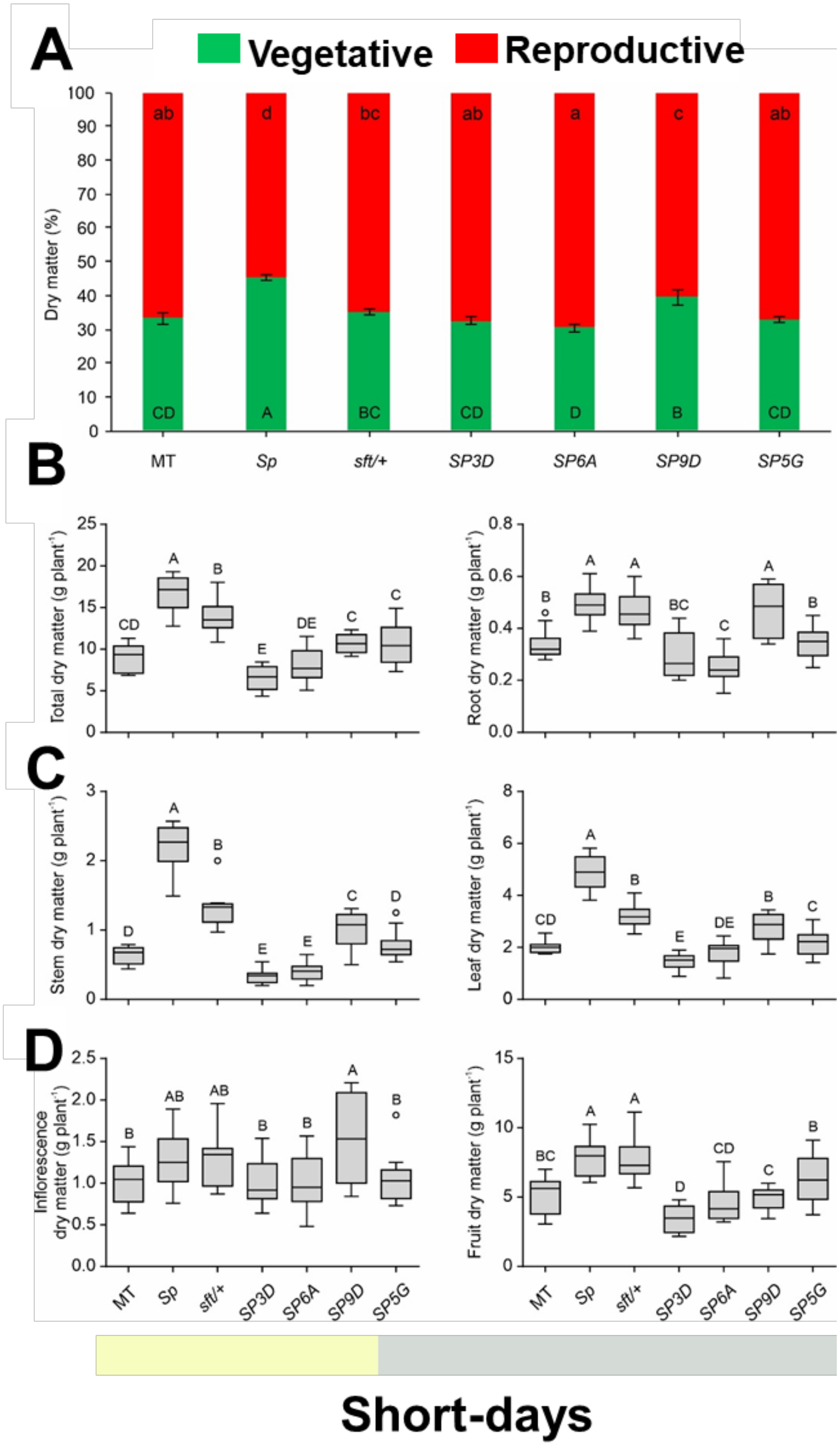


**Supplementary Figure S8. Vegetative to reproductive balance on tomato genotypes carrying distinct alleles from the *SELF-PRUNING* gene family under short-days. A**, dry matter (%) distribution between vegetative (roots, stem and leaves) and reproductive (inflorescences, flowers and fruits) organs. **B**, total and root dry matter of plants. **C**, shoot and leaf dry matter of plants. **D**, inflorescence and fruit dry matter of plants. The values were obtained from represent 8-12 plants per genotype, and statistical groups were determined using a Tukey honest significant difference (HSD) test ($P$ <0.05) and are indicated with different letters.

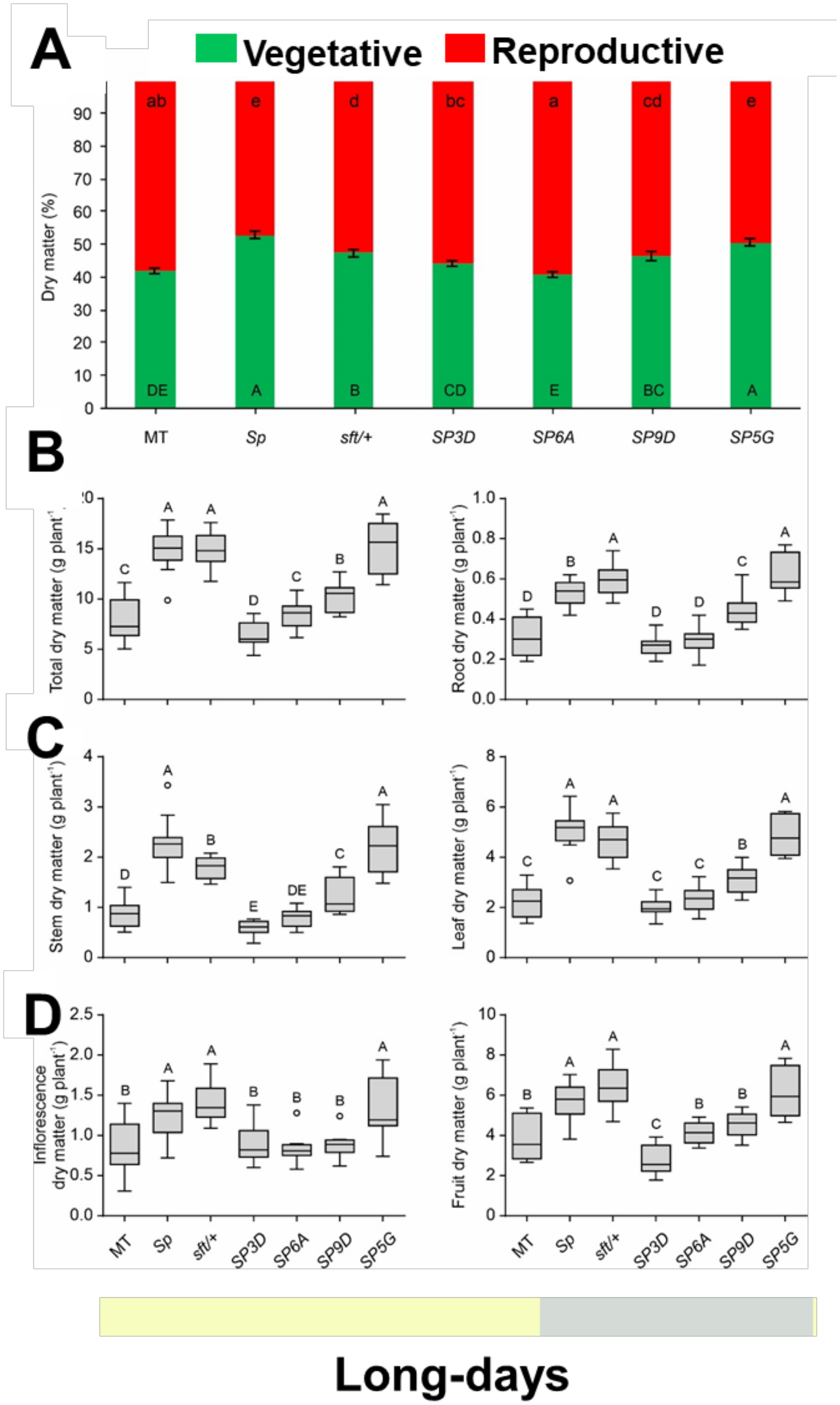


**Supplementary Figure S9. Vegetative to reproductive balance on tomato genotypes carrying distinct alleles from the SELF-PRUNING gene family under long-days**. **A**, dry matter (%) distribution between vegetative (roots, stem and leaves) and reproductive (inflorescences, flowers and fruits) organs. **B**, total and root dry matter of plants. **C**, shoot and leaf dry matter of plants. **D**, inflorescence and fruit dry matter of plants. The values were obtained from represents 8-12 plants per genotype, and statistical groups were determined using a Tukey honest significant difference (HSD) test (*P* <0.05) and are indicated with different letters.

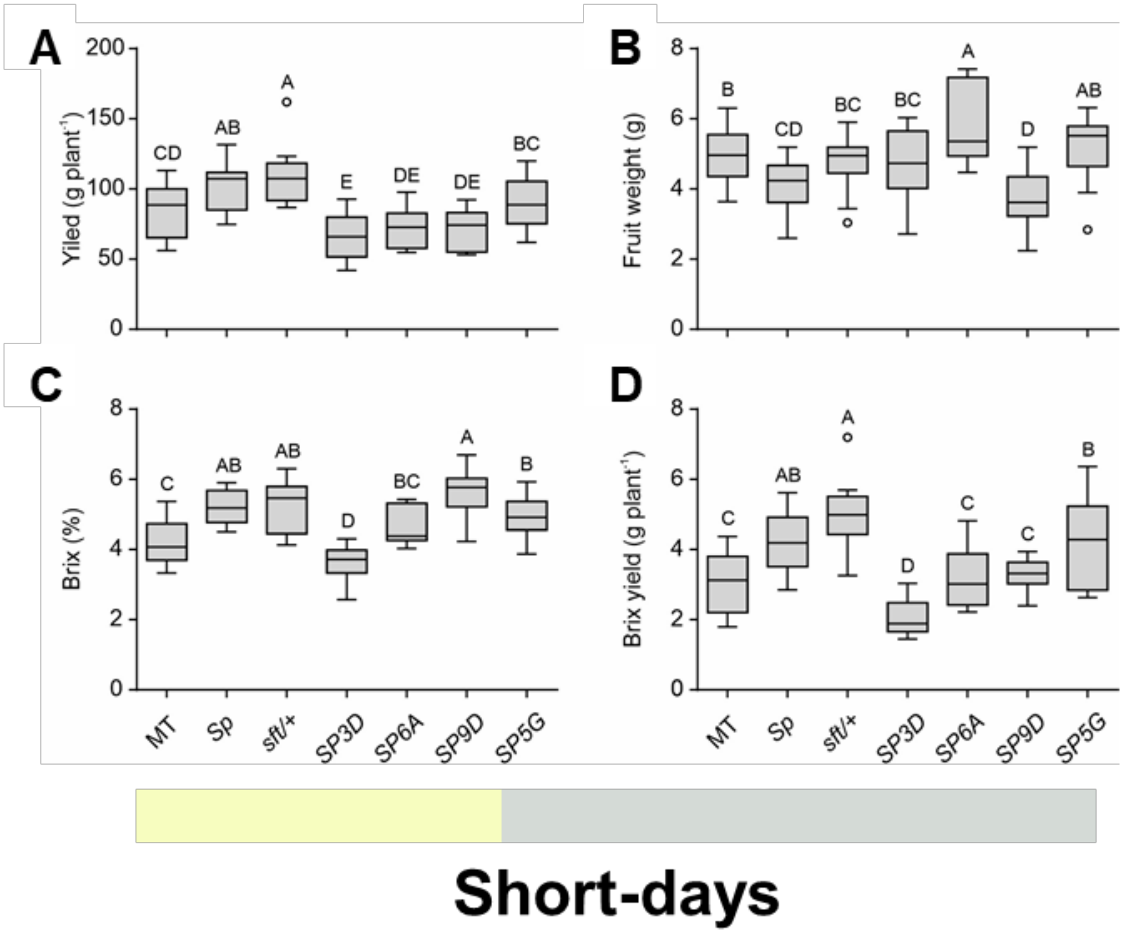


**Supplementary Figure S10. Tomato productivity traits under short-days in tomato genotypes carrying distinct alleles from the *SELF-PRUNING* gene family. A**, fruit production per plant (yield). **B**, mature fruit weight. **C**, percentage of total soluble solids for 30 fruits (Brix (%)). **D**, relationship between Brix (%) and mature fruits at 127 days after germination. The data were obtained from 8-12 plants per genotype, and statistical groups were determined using a Tukey honest significant difference (HSD) test (*P* <0.05) and are indicated with different letters.

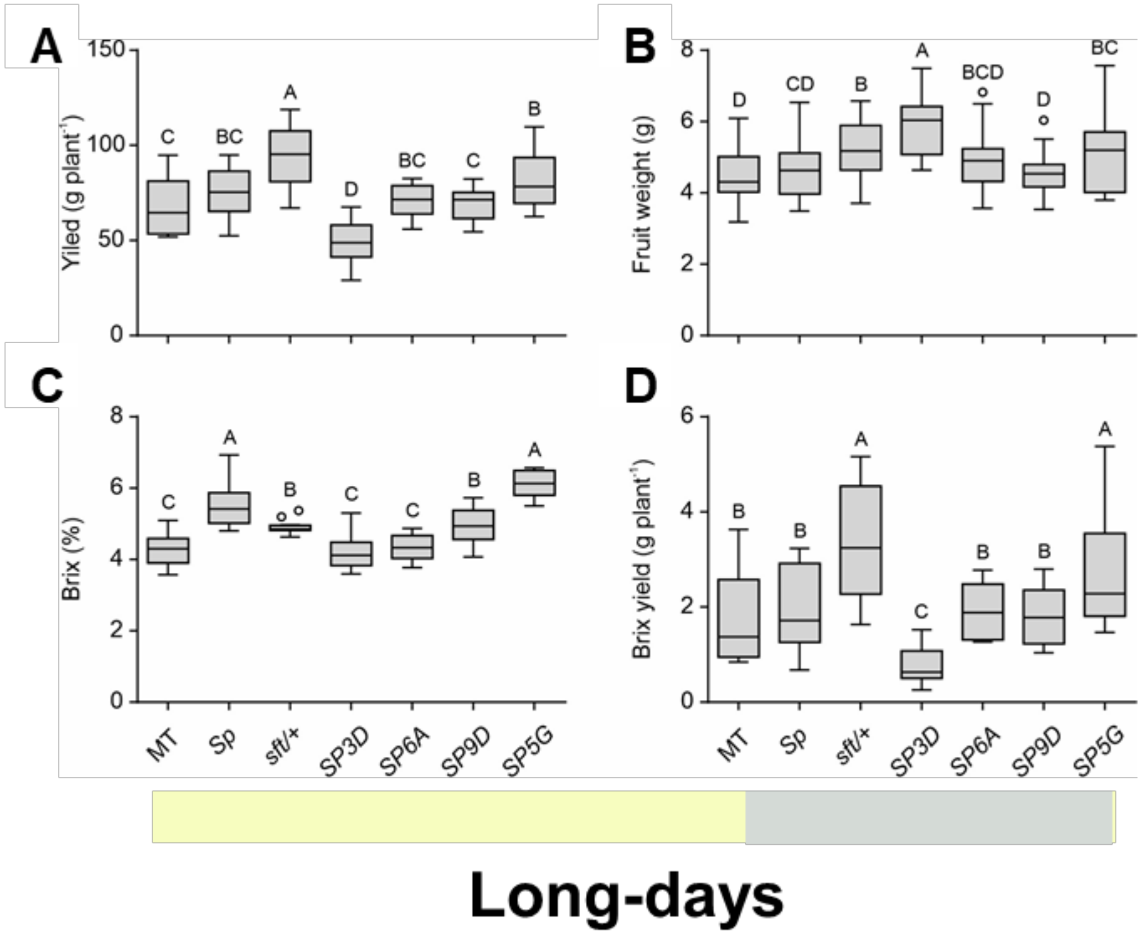


**Supplementary Figure S11. Tomato productivity traits under long-days in tomato genotypes carrying distinct alleles from the SELF-PRUNING gene family. A**, fruit production per plant (yield). **B**, mature fruit weight. **C**, percentage of total soluble solids for 30 fruits (Brix (%)). **D**, relationship between Brix (%) and mature fruits at 127 days after germination. The data were obtained from 8-12 plants per genotype, and statistical groups were determined using a Tukey honest significant difference (HSD) test (*P* <0.05) and are indicated with different letters.

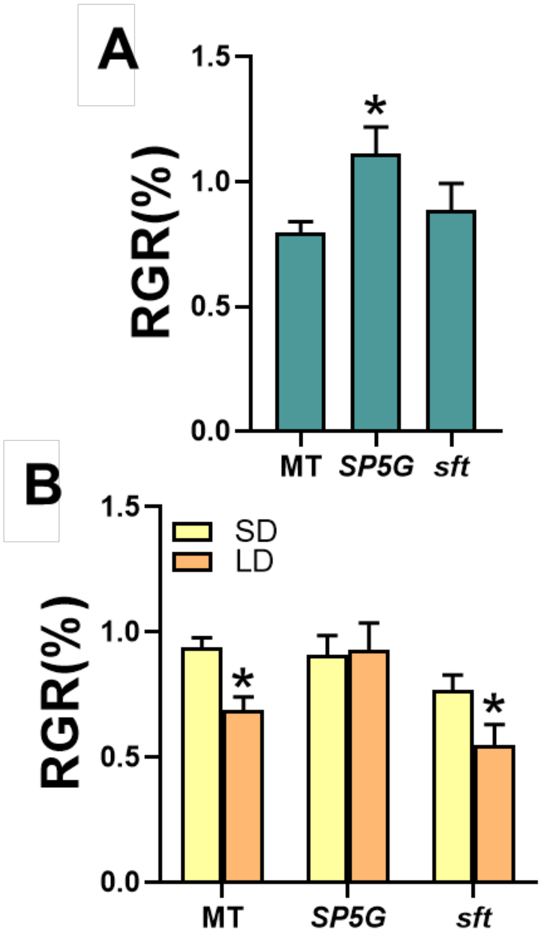


**Supplementary Figure S12. Tomato photoperiod responses modulate root elongation and aluminium (Al) tolerance. A**, Relative growth rate (RGR - %) was determined by the root elongation ratio measured in plants growing at short-days (SD) or long-days (LD) under optimal control conditions. Root elongation was determined in 5-day-old seedlings cultivated on hydroponics culture. RGR (%) was calculated as followed: SD root elongation (cm) / LD root elongation (cm) (n = 12 plants for each condition). An asterisk (*) indicate mean values that were determined by the two-sided Student's *t*-test to be different ($P < 0.05$) from the wild-type tomato Micro-Tom (MT). **B**, RGR (%) indicated root elongation measured in plants growing in the absence of $Al^{3+}$ or in the presence of 100 µM $Al^{3+}$ under either SD or LD. RGR (%) was calculated as follow: SD (-Al) root elongation (cm) / SD (+Al) root elongation (cm) (yellow bars); and LD (-Al) root elongation (cm) / LD (+Al) root elongation (cm) (orange bars). An asterisk (*) indicate values that were determined by the two-sided Student's *t*-test to be different ($P < 0.05$) between growth conditions (SD and LD). Abreviations: MT, *Solanum lycopersicum* cv. Micro-Tom; *SP5G*, *SELF-PRUNING 5G* allele from *Solanum pennellii* introgressed into MT; *SFT*, introgression of a loss-of-function mutation on *SINGLE FLOWER TRUSS.*

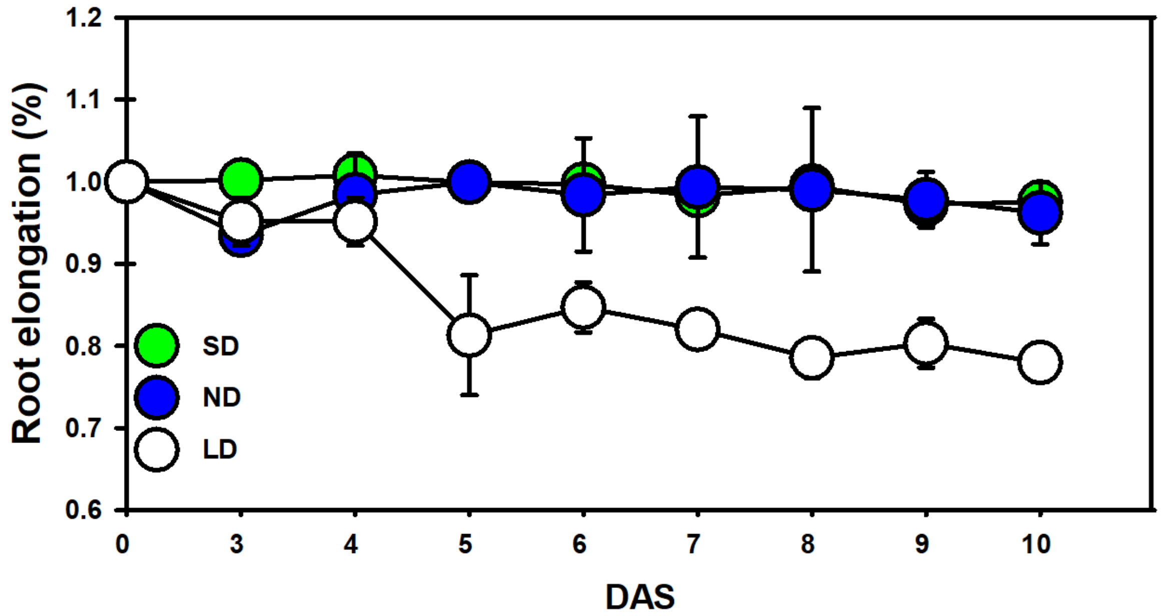


**Supplementary Figure S13. Root elongation under aluminium (Al) stress is associated with day-length.** Root elongation (%) in *Arabidopsis thaliana* ecotype Columbia (Col-0) was determined during 10days in seedlings growing at pH 4.0 in the absence of $Al^{3+}$ (-Al) or in the presence of 50 µM $Al^{3+}$ (+Al) under at either short-day (SD), neutral-day (ND), or long-day (LD) conditions (n = 60 from three independent experiments). Root elongation (%): -Al-root elongation (cm) / +Al-root elongation (cm).

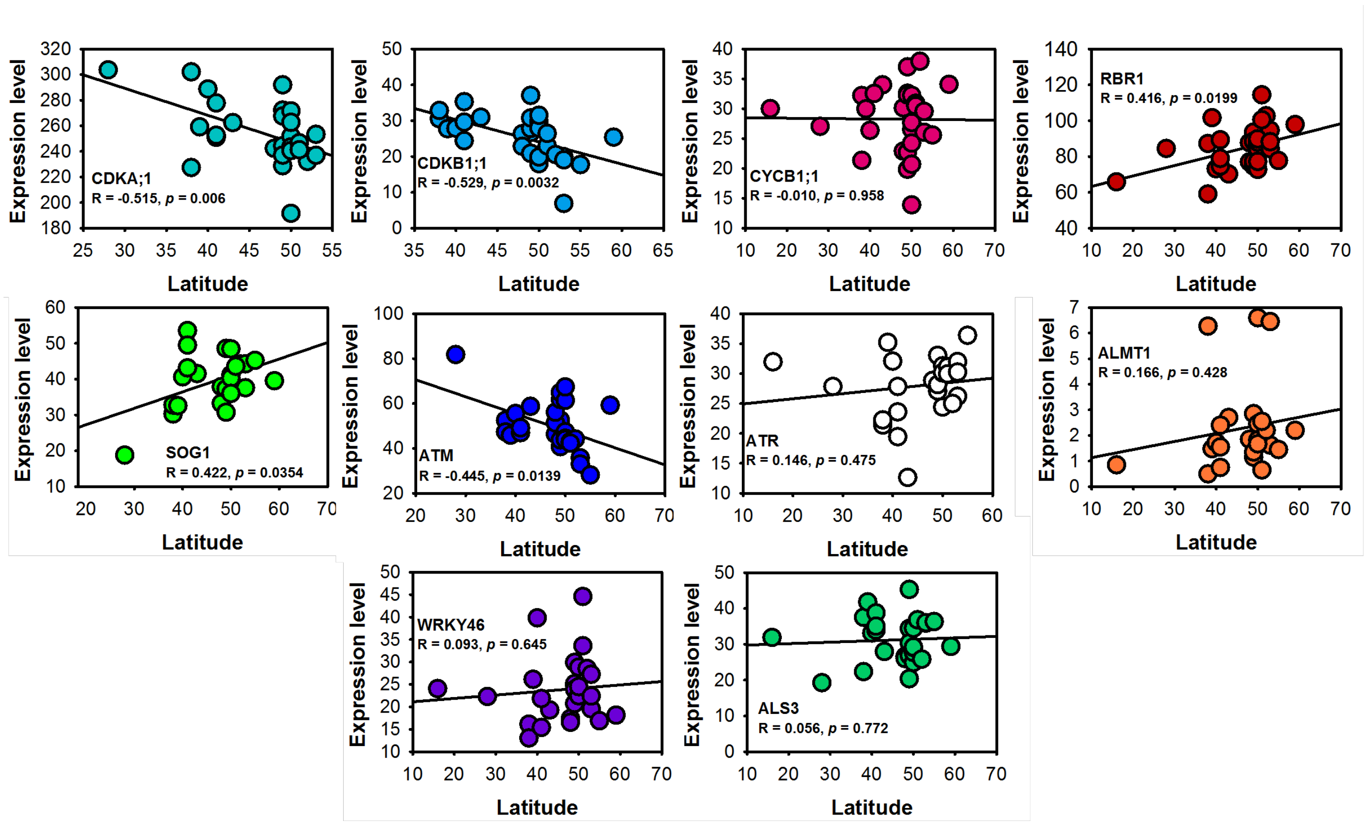


**Supplementary Figure S14. Pearson Correlations between *in silico* gene expression and latitudinal variation**. Data represent analyses using 30 *Arabidopsis thaliana* accessions with centre of origin from contrasting latitude (http://bar.utoronto.ca/). Abbreviations: *CDKA*;1 (Cyclin-*dependent* kinase *A;1*); *CDKB1;1* (Cyclin-*dependent* kinase *B1;1*); *CYCB1;1* (*Cyclin B1;1*); *Retinoblastoma-related 1* (*RBR1*); *SOG1* (*SUPPRESSOR OF GAMMA RESPONSE1*), *ATM* (*ATAXIA TELANGIECTASIA MUTATED*); *ATR* (*ATAXIA TELANGIECTASIA AND RAD3 RELATED*); *ALMT1* (*Aluminum-activated malate transporter 1*); *WRKY46* (*WRKY transcription factor 46*); *ALS3* (*ALUMINUM SENSITIVE 3*).

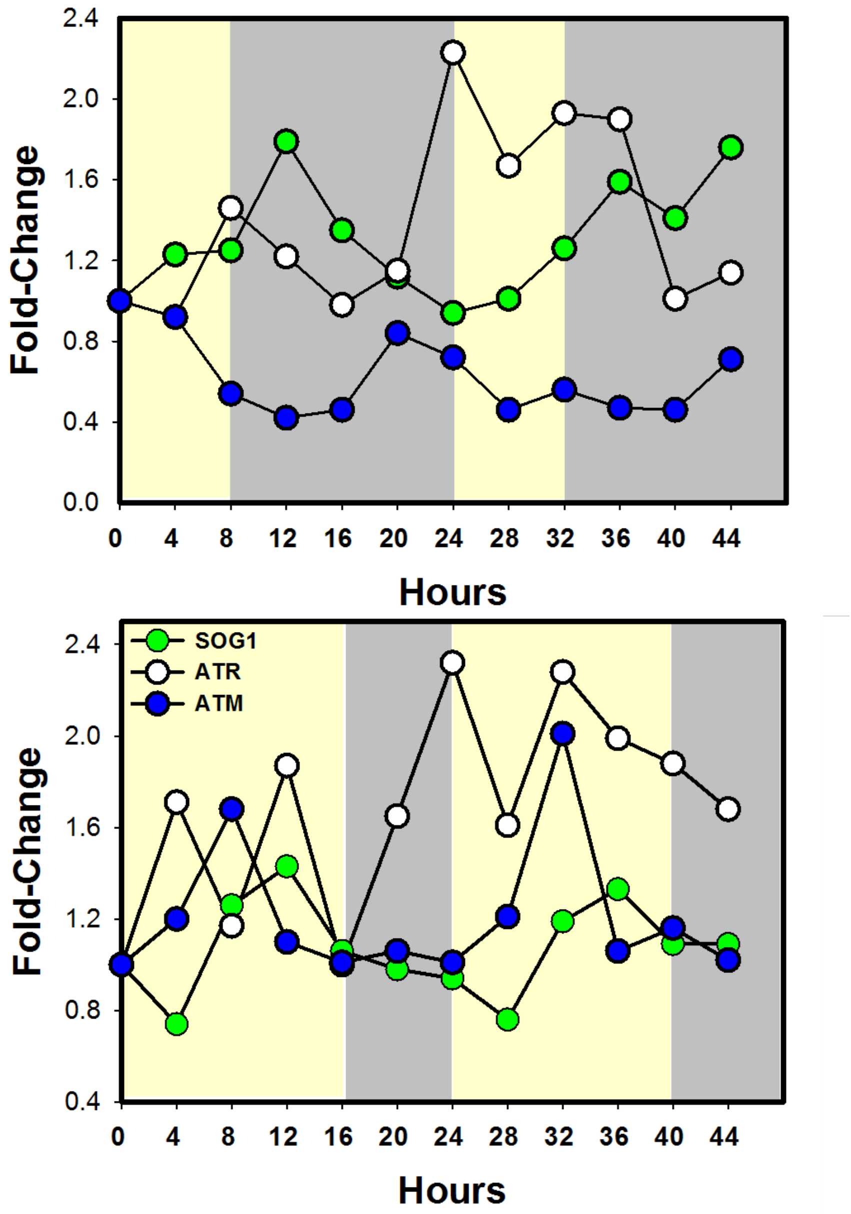


**Supplementary Figure S15. *In silico* gene expression for regulators of DNA checkpoints under different photoperiod length.** *In silico* transcripts profile for *SOG1* (*SUPPRESSOR OF GAMMA RESPONSE1*), *ATM* (*ATAXIA TELANGIECTASIA MUTATED*), and *ATR* (*ATAXIA TELANGIECTASIA AND RAD3 RELATED*) were obtained using the publicly data available at the (http://bar.utoronto.ca/). Light and dark rectangles denotes day and night periods under short-days (SD) and long-days (LD), respectively.

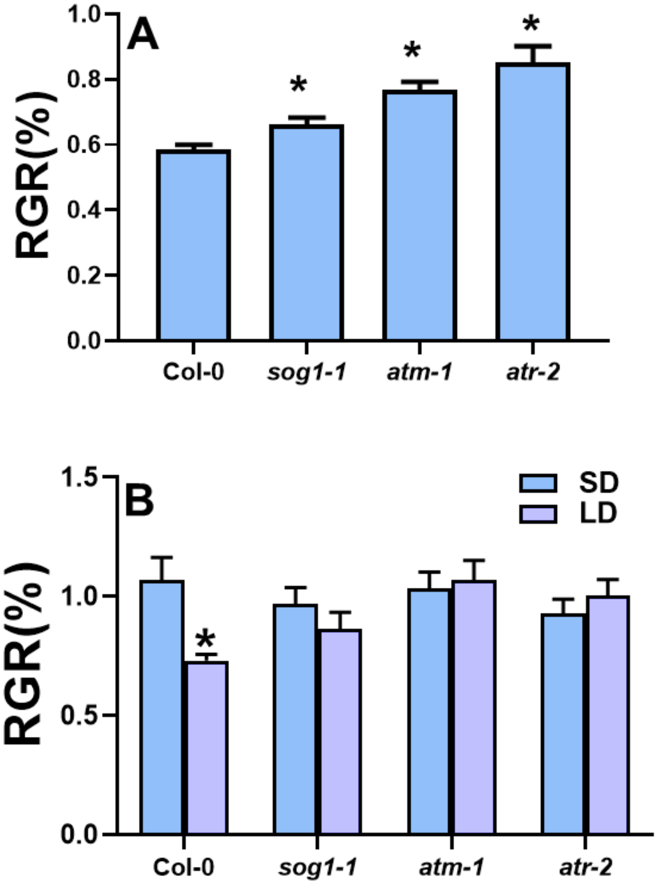


**Supplementary Figure S16. DNA checkpoint and endoreduplication related genes are implicated on aluminium (Al) tolerance modulated by day-length. A**, Relative growth rate (RGR - %) was determined by the root elongation ratio measured in plants growing at either short-days (SD) or long-days (LD) under control conditions (absence of Al). Root elongation was determined in 10-day-old seedlings cultivated on MS medium. RGR (%) was calculated as followed: SD root elongation (cm) / LD root elongation (cm) (n = 40 plants for each condition). An asterisk (*) indicate values that were determined by the two-sided Student's *t*-test to be different ($P < 0.05$) to the *Arabidopsis thaliana* wild type ecotype Columbia-0 (Col-0). **B**, RGR (%) describes measurements of the root elongation in wild-type and *A. thaliana* mutants plants for DNA-repair and endoreduplication related genes growing in the absence of $Al^{3+}$ or in the presence of 50 μM $Al^{3+}$ under either SD or LD. RGR (%) was calculated as follow: SD (-Al) root elongation (cm) / SD (+Al) root elongation (cm) (blue bars); and LD (-Al) root elongation (cm) / LD (+Al) root elongation (cm) (orange bars) (n = 40 plants for each condition). An asterisk (*) indicate values that were determined by the two-sided

Student's *t*-test to be different ($P < 0.05$) between SD and LD. Abbreviations: *sog1-1* (*Suppressor of Gamma Response1-1*), *atm1* (*Ataxia T Telangiectasia Mutated-1*), *atr-2* (*Ataxia Telangiectasia and Rad3 Related-2*).

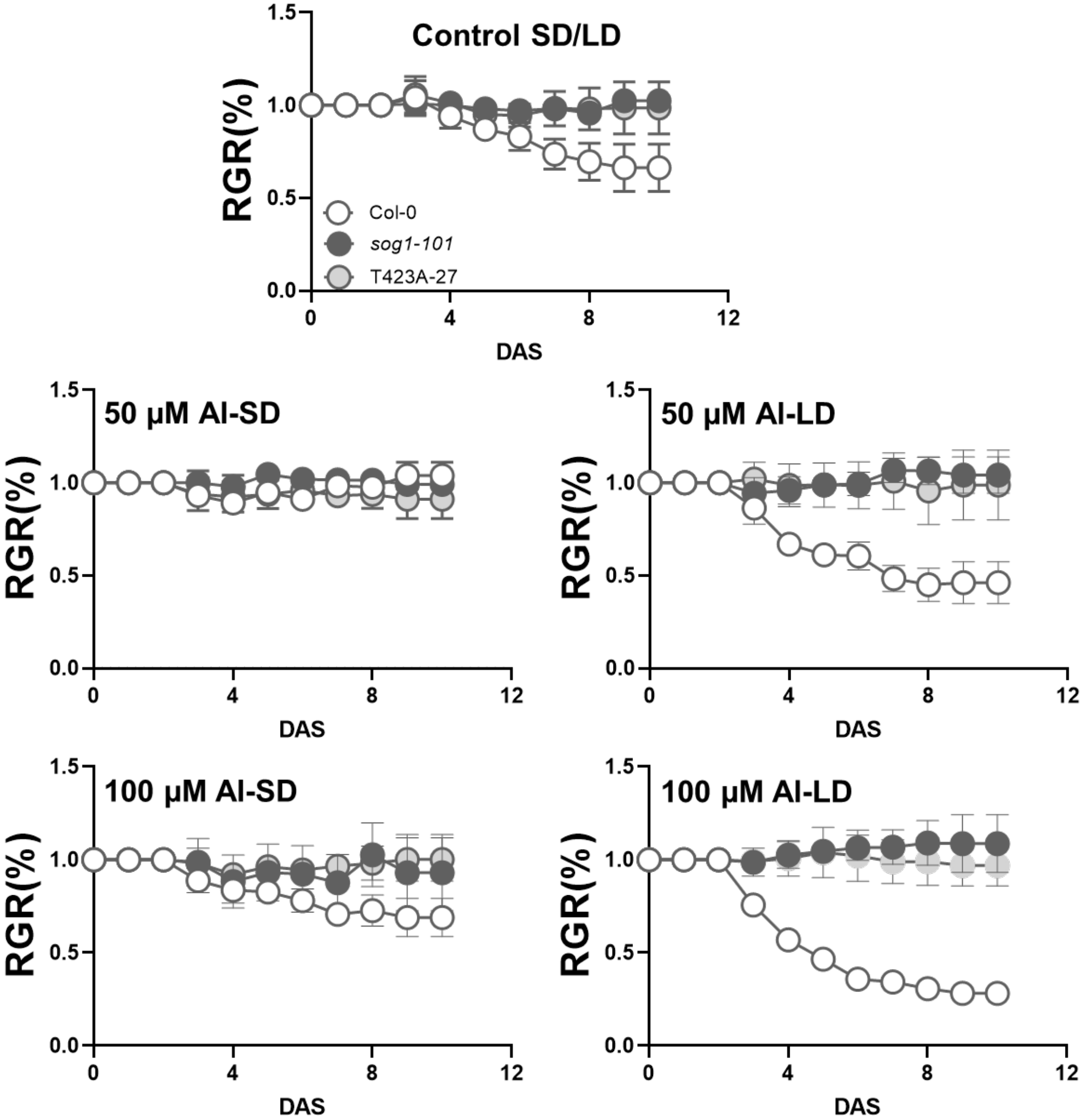


**Supplementary Figure S17. Aluminium tolerance is dependent of the circadian clock related protein casein kinase 2 (CK2).** Wild-type (Col-0), s*og1* knockout mutant *sog1-101* and phospho-mutant (*T423A-27*) on *sog1* gene, site phosphorylated by CK2. Relative growth rate (RGR -%) was determined in seedlings with 10-old-days growing at pH 4.0 in the absence of $AlCl_3$ or in the presence of 50 or 100 µM $AlCl_3$ under either short-days (SD) or long-days (LD) (n = 5, with 6 plants in each repeat) during 10 days after showing (DAS). RGR (%): -Al-root elongation (cm) / +Al-root elongation (cm) Abbreviations: *sog1-1* (*Suppressor of Gamma Response1-1*); SD, short-days; LD, long-days; DAS, days after sowing.

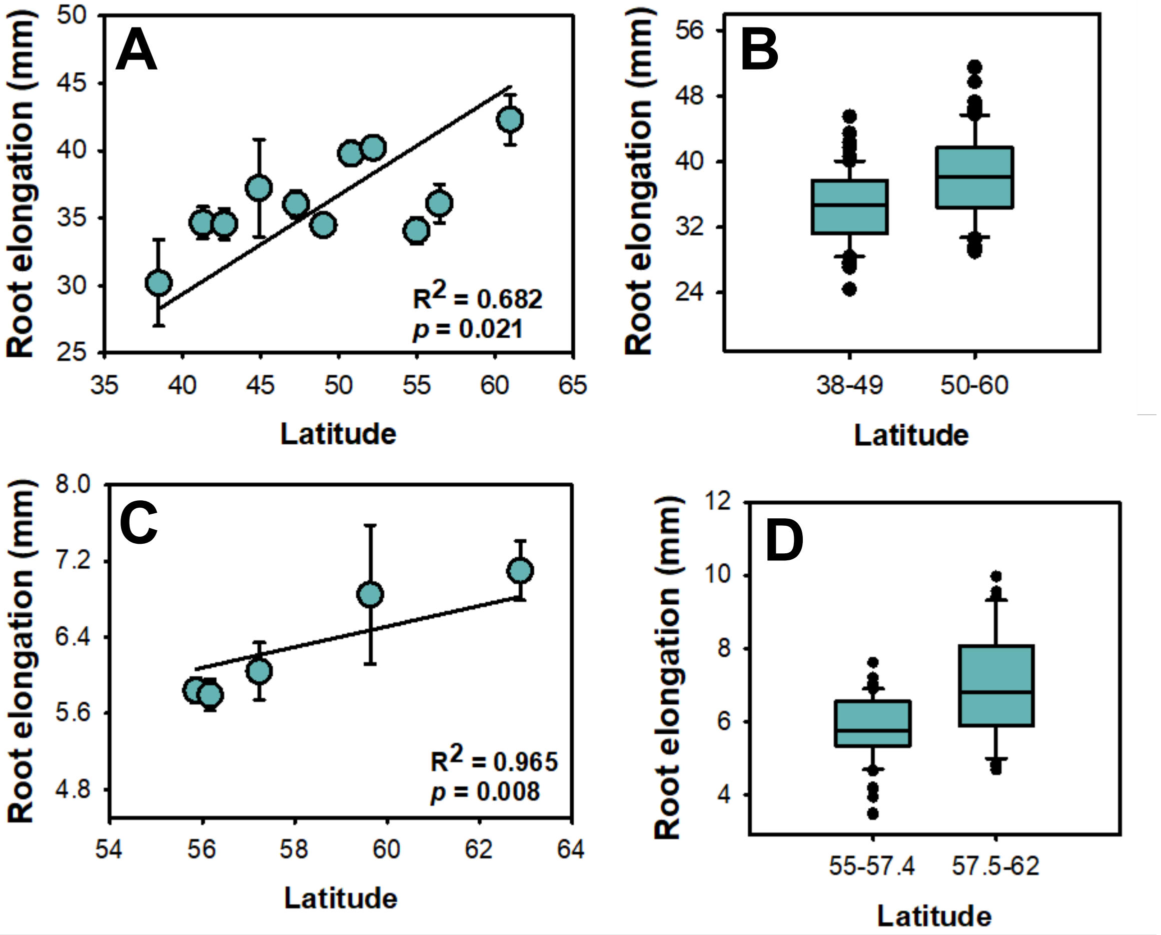


**Supplementary Figure S18. Relationships between root elongation and latitudinal variation in *Arabidopsis thaliana* accessions. A-B**, represent data from Ristova *et al.* (2018). **C-D**, represent data from Satbhai *et al.* (2017). **A**, **C** exhibit Pearson correlations between latitude and root elongation for groups obtained in the AraPheno database (https://arapheno.1001genomes.org/). **B**, **D** demonstrate the comparison between groups from two latitudinal ranges.

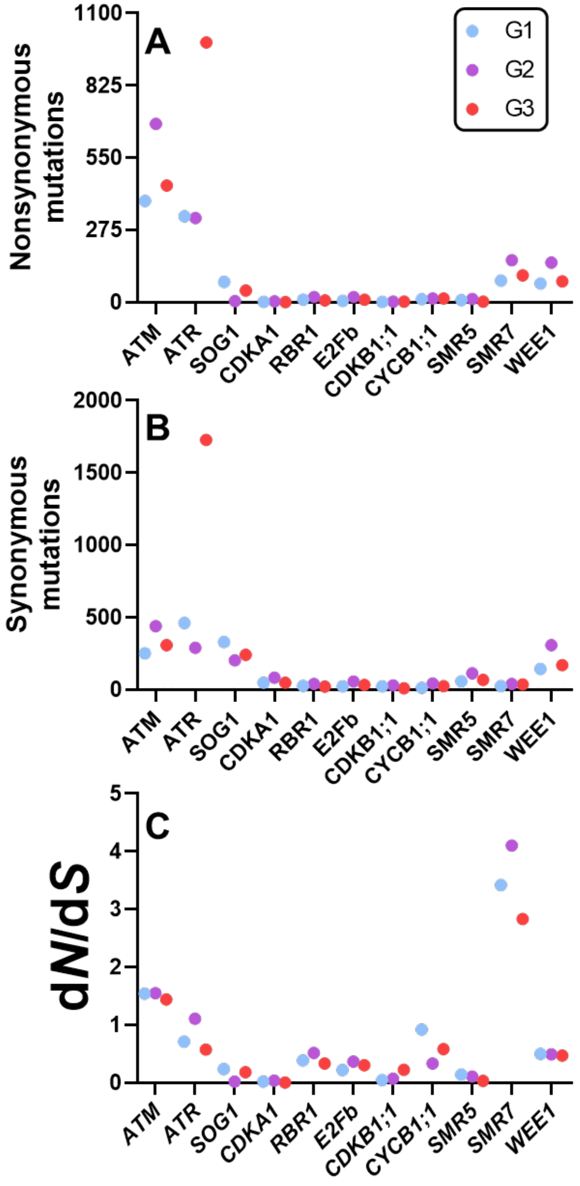


**Supplementary Figure S19. Distribution of nonsynonymous, synonymous, and ratio between nonsynonymous and synonymous fractions.** 287 *Arabidopsis thaliana* accessions were separated by grouping analyses, G1 (short-days), G2 (neutral), and G3 (long-days), and were further submitted to the POLYMORPH 1001 (https://tools.1001genomes.org/polymorph/) to check the existence of genetic variability on genes involved on DNA repair, cell cycle, and endoreduplication. Polymorphisms revealed the presence of (**A**) nonsynonymous, and (**B**) synonymous mutations on the aforementioned genes as well as the ratio of nonsynonymous to synonymous fractions (*dN*/*dS*) (**C**). The genes investigated were *ATM* (*Ataxia T Telangiectasia Mutated*), *ATR* (*Ataxia Telangiectasia and Rad3 Related-2*), *SOG1* (*Suppressor of Gamma Response1-1*), *CDKA1* (*Cyclin-dependent kinase A1*), RBR1 (*Retinoblastoma-related 1*), *E2Fb* (E2F transcription Factor B), *CDKB1;1* (*Cyclin-dependent kinase B1;1*), *CYCB1;1* (*Cyclin B1;1*), *SIAMESE-RELATED 5* (*SMR5*), *SIAMESE/SIAMESE-RELATED 7* (*SMR7*) and *wee1-1* (Mitosis inhibitor protein kinase wee1).

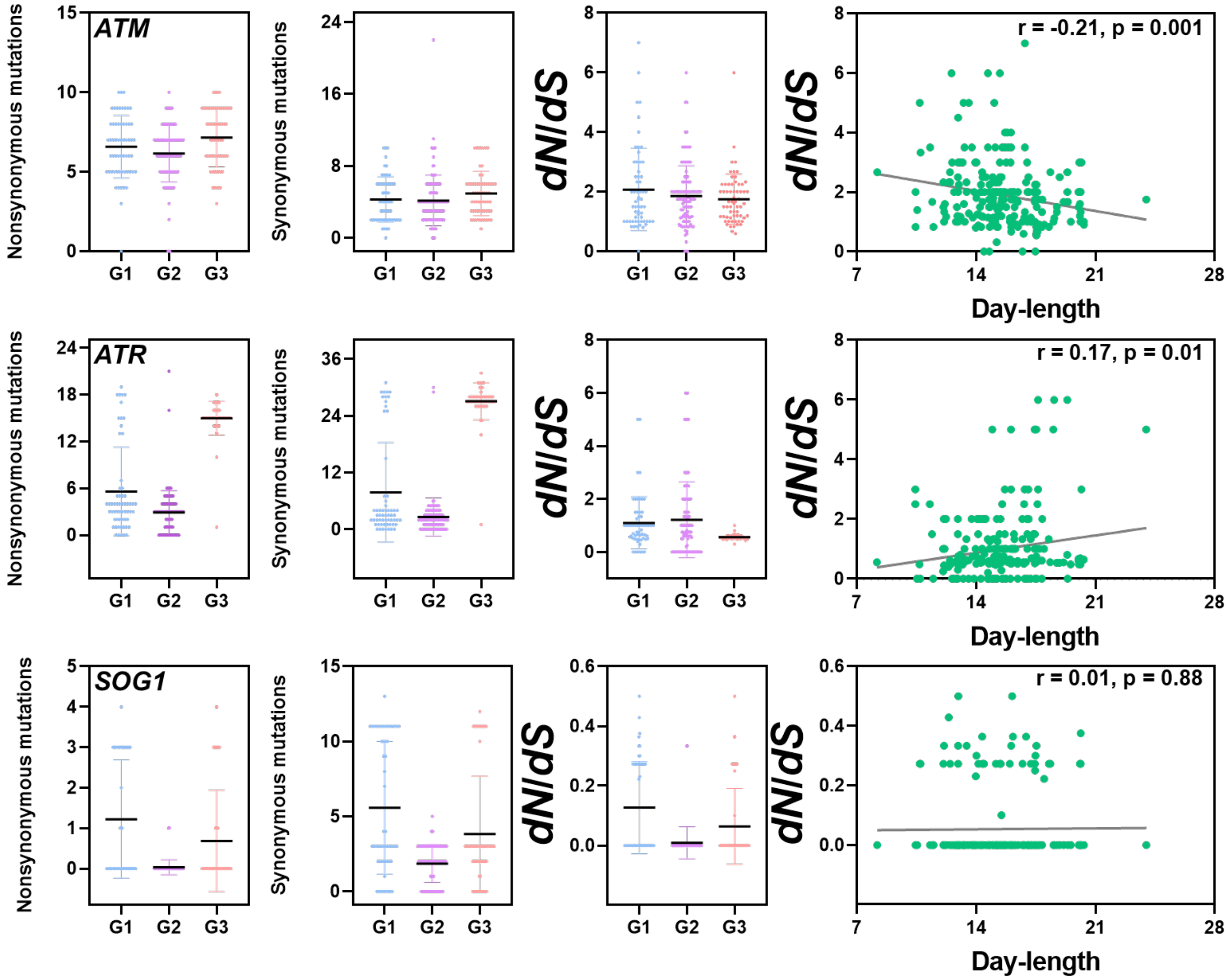


**Supplementary Figure S20. Comparison of nonsynonymous, synonymous and ratio of nonsynonymous to synonymous fractions between *Arabidopsis thaliana* groups.** By using the POLYMORPH 1001 (https://tools.1001genomes.org/polymorph/), we compared polymorphisms occurring on the genes *ATAXIA TELANGIECTASIA MUTATED* (*ATM*), *ATAXIA TELANGIECTASIA AND RAD3 RELATED* (*ATR*), and *SUPPRESSOR OF GAMMA RESPONSE1* (*SOG1*) among groups G1, G2, and G3, in which 287 *A. thaliana* accessions were separated. Additionally, the ratio of nonsynonymous to synonymous fractions (*dN*/*dS*) was further correlated with day-length (hours).

**Supplementary Table S2.** List of plant material used in this study.

| Specie | Genotype | Gene ID | Mutation | References |
|---|---|---|---|---|
| *A. thaliana* | Columbia-0 (Col-0) | - | Wild-Type | |
| *A. thaliana* | *atm-1* | AT3G48190 | An insertion in the gene 3' end is found in this line, representing alternative allele. The T-DNA insertion is on exon 78 resulting in a 23-bp deletion at ATM sequence, and their background is Wassilewskija. | Garcia et al (2003) |
| *A. thaliana* | *atr-2* | AT5G40820 (SALK_032841) | The T-DNA is on ATR gene middle in fraction corresponding to exon 10, representing the Columbia background. | Culligan et al. 2004 |
| *A. thaliana* | *sog1-1* | AT1G25580 | A missense mutation occurs in the highly conserved amino acid region at NAC domain, wherein Gly (GGA)-155 is changed to Arg (AGA). | Yoshiyama et al. (2009) |
| *A. thaliana* | *sog1-101* | AT1G25580 (GABI_602B10) | Homozygous knockout mutant line. | Ogita et al. (2018) |
| *A. thaliana* | *sog1-1* (*T423A-27*) | AT1G25580 | The threonine positioned at 423 amino acid was replaced to alanine (*T423A*). The resulting allele was used to complement knockout line *sog1-101*. | Wei et al. (2020) |
| *A. thaliana* | *e2fb-1* | AT5G22220 (SALK_103138) | A T-DNA insertion representing a knockout gene line, showing the inserted sequence in the exon 9 that abolish the gene expression. | Berckmans et al. (2011) |
| *A. thaliana* | *CDKB1* | AT3G54180 | The codon GAT (Asp-161) was changed to AAT (Asn-161) by using *in vitro* mutagenesis, resulting the dominant negative *CDKB1;1.N161* allele. This allele was expressed under control of the constitutive 35S promoter of the Cauliflower mosaic virus (CaMV35S). The amino acid residue at position 161-Asp is required for correct ATP binding by CDK kinases, where the mutation compromises the kinase activity. | Boudolf et al. (2004) |
| *A. thaliana* | *smr5* | AT1G07500 (SALK_100918) | Line with a T-DNA insertion on the position 2304967bp of chromosome 1, representing a knockout line for the gene *SMR5.* | Yi et al. (2014 |
| *A. thaliana* | *smr7* | AT3G27630 (SALK_128496) | Line with a T-DNA insertion on the position 10231203bp at chromosome 3, representing a knockout line for the gene SMR7. | Yi et al. (2014 |
| *A. thaliana* | *wee1-1* | AT1G02970 | The insertion of T-DNA is located on intron between exons 7 and 8, which result in a deletion of the last 197 amino acids, representing probably null alleles. | Schutter et al. (2007) |
| *A. thaliana* | *wee1-2* | AT1G02970 | A T-DNA insertion on the exon 1, deleting most of the protein, also a probable null allele. | Schutter et al. (2007) |
| *S. lycopersicum* | MT | - | A cultivar containing the recessive allele self-prunning (*sp*) making the growth habit determinated and turning fruit ripening uniform. This transcription factor is at chromosome 6, and acts repressing the flowering. | - |

| | | | | |
|---|---|---|---|---|
| *S. lycopersicum* | *Sp* | Solyc06g074350 | Resulting from an introgression of allele *Self-pruning* (*Sp*) from cv. Moneymaker on MT cultivar. This result in a resumed vegetative growth after flower induction, which maintain the vegetative phase on plant. | Tomato esalq.usp seed-bank |
| *S. lycopersicum* | *sft* | Solyc03g063100 | The MT cultivar was in introgressed with the loss-of-function mutation on single flower truss (*sft*) (LA2460) originating near-isogenic lines. The mutation promotes low flowering induction, and the transcription is at chromosome 3. | Tomato esalq.usp seed-bank |
| *S. lycopersicum* | SP5G | Solyc05g053850 | The MT cultivar was in introgressed with the function allele of flowering repressor SELF-PRUNING 5G (SP5G) from *Solanum pennellii* (LA2460) originating near-isogenic lines. The allele repress the flowering under long-day conditions promotes low flowering induction, and the gene is located at chromosome 5. | Tomato esalq.usp seed-bank |
| *S. lycopersicum* | M82 | | Wild-type | - |
| *C. juncea* | IAC-KR 1 | - | Wild-Type | - |
| *P. sativum* | Amélia | - | Wild-Type | - |
| *L. culinaris* | Silvina | - | Wild-Type | - |
| *S. humilis* | BGH-UFV | - | Wild-Type | - |
| *V. unguiculata* | Pingo de Ouro | - | Wild-Type | - |
| *L. albus* | TRM 881 | - | Wild-Type | - |

**Supplementary Table S3.** Primer sequences used in qRT-PCR analyses performed in this study

| Gene | Locus | Forward | Reverse |
|---|---|---|---|
| RBR1 | AT3G12280 | 5'-GTGATCACCTCAGGCTATGAGC-3' | 5'-CACTCTTTCTAACTGTGTTCGGTGT-3' |
| SOG1 | AT1G25580 | 5'-CCATGAGGTTTCTCTTGCCGAGAC-3' | 5'-TCAGGCCCAGAACTTGGTCTTTC-3' |
| ATR | AT5G40820 | 5'-GTGCCATTCAGATTGACCCAGAAC-3' | 5'-TGCCCTCATATCCAGTGATGCC-3' |
| ATM | AT3G48190 | 5'- AACTTGATGGCTACGAGGGTGGT -3' | 5'- CCAGGGAACATATGGGACAAACG -3' |
| CYCB1 | AT4G37490 | 5'-ACCTCGCAGCTGTGGAATATGTG-3' | 5'-ATCTCGTGGCCTCCATTCACTCTC -3' |
| CDKA1 | AT3G48750 | 5'-ACTGGCCAGAGCATTCGGTATC -3' | 5'-TCGGTACCAGAGAGTAACAACCTC-3' |
| CDKB1 | AT3G54180 | 5'- ACTGGTGTTGACATGTGGTCTG -3' | 5'- TTGGTGTTCCTAGCAACCTGAAG -3' |

**Additional files**

**Data S1. Number of principal component analyses (PCA) retained:** a−score optimisation − spline interpolation.

**Data S2. Principal component analyses (PCA):** Variance explained by PCA.

**Data S3. Pop_Orange_Arabidopsis:** Grouping analysis separating *Arabidopsis thaliana* accessions according with Orange Canvas software, for details see material and methods.

**Data S4. Scatter plot:** Representation of three group distinction.

**Data S5. Barplot:** % of reassignment to actual group.

**Data S6. Compoplot:** membership probability for three groups.

**File S1.** ***Arabidopsis thaliana*** **accessions details**
