## Supplementary figures and images for "Photoperiod shapes aluminium tolerance in plants"

### Data S1

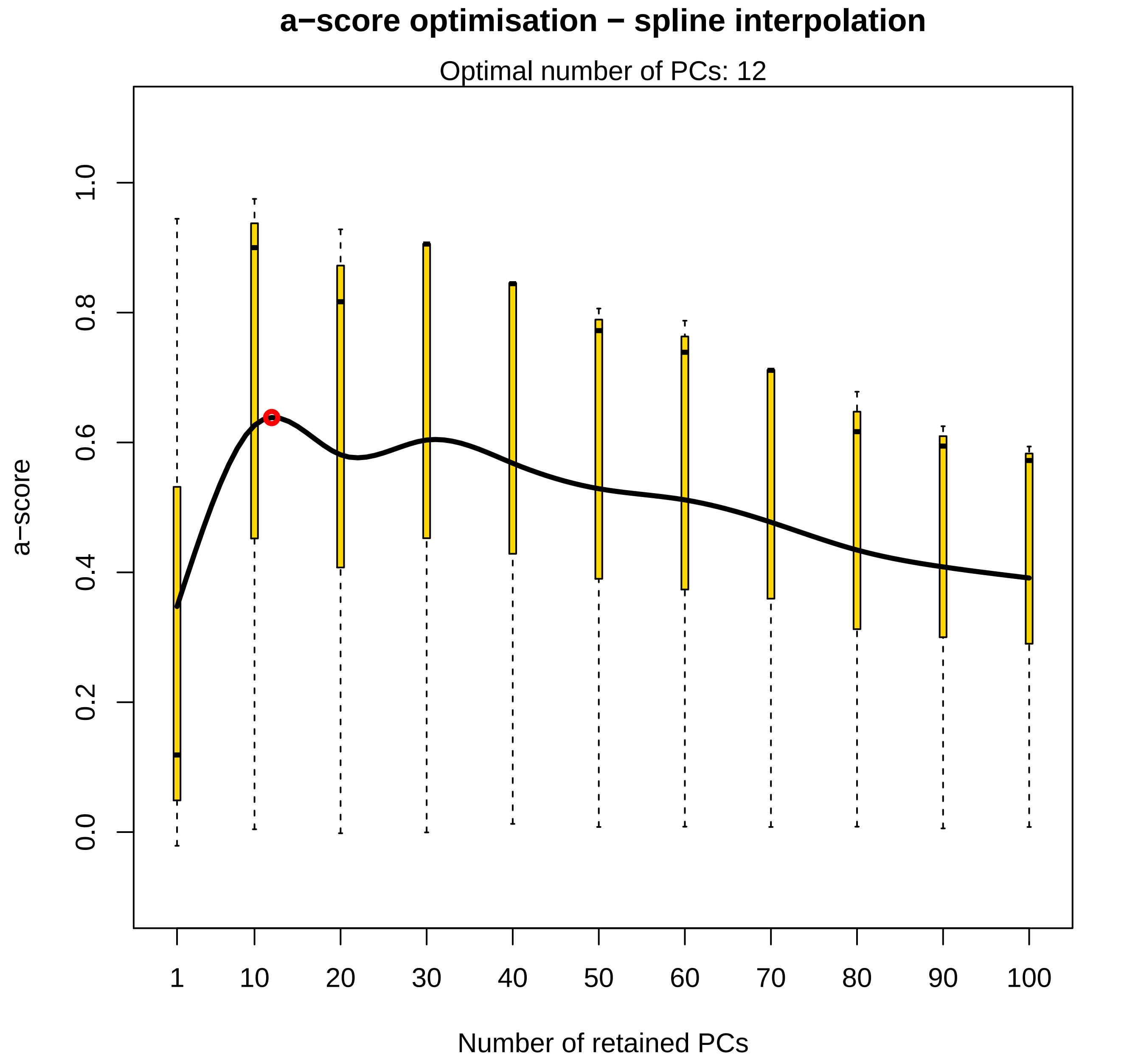

### Data S2

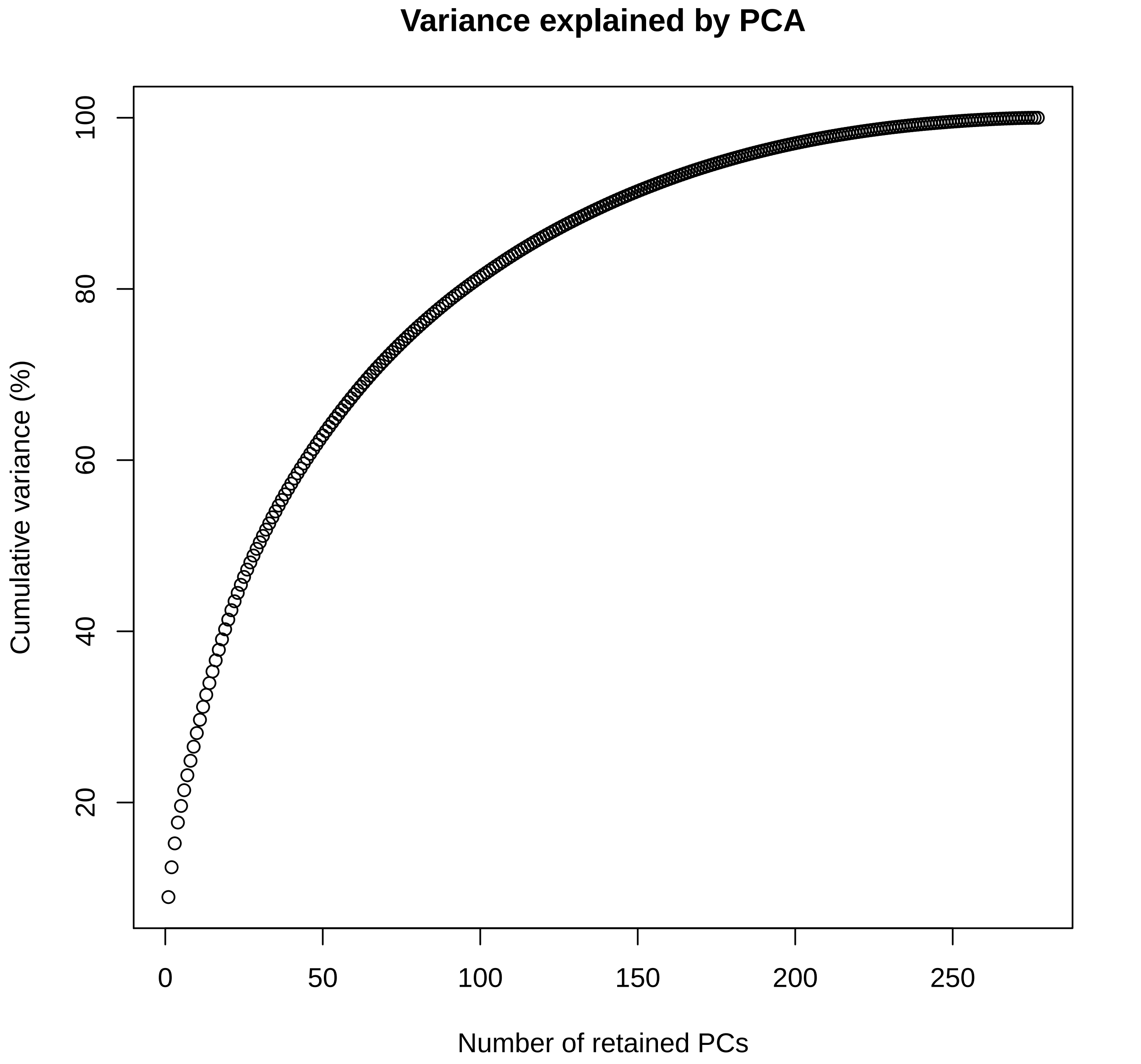

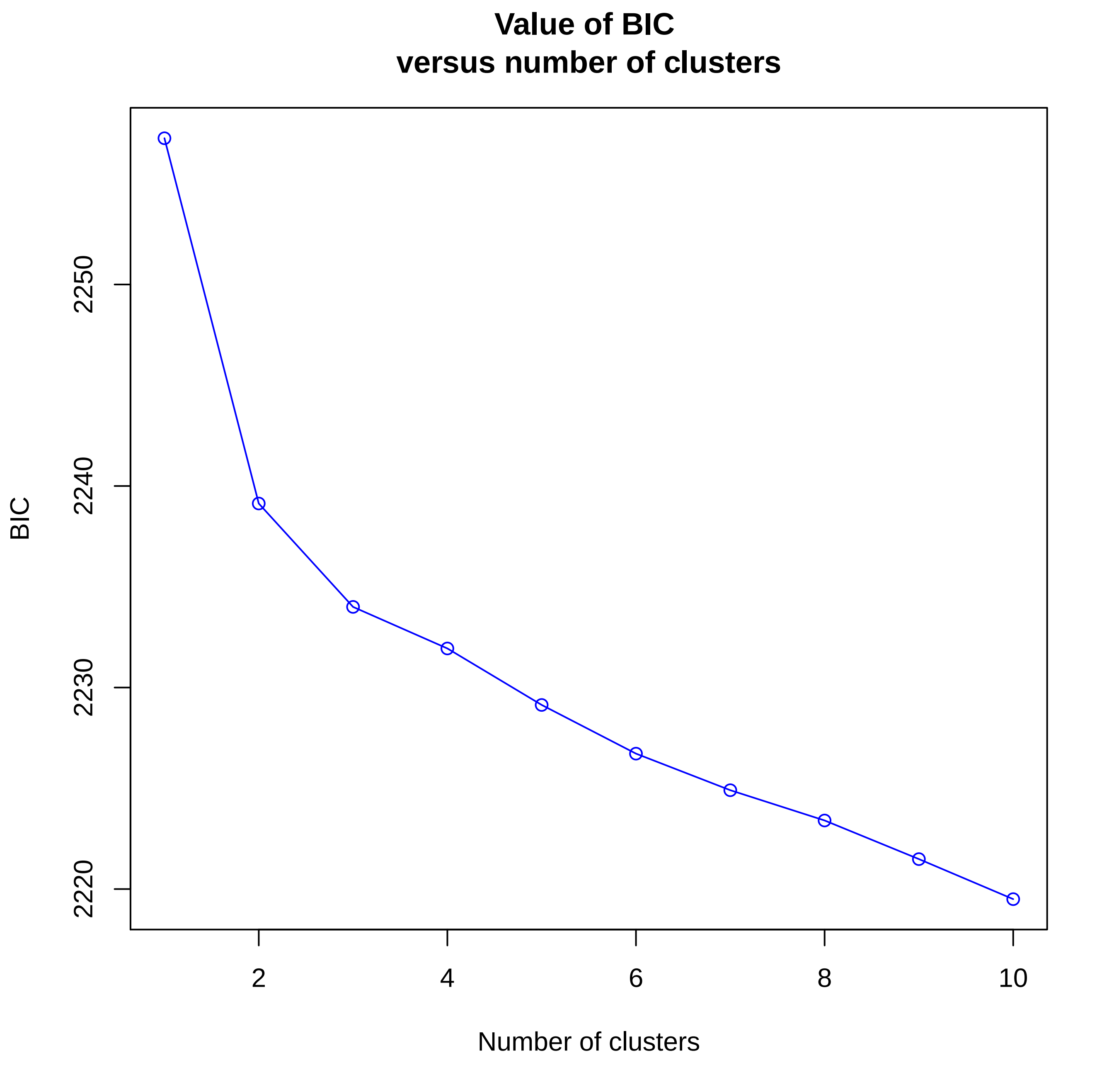

### Data S3

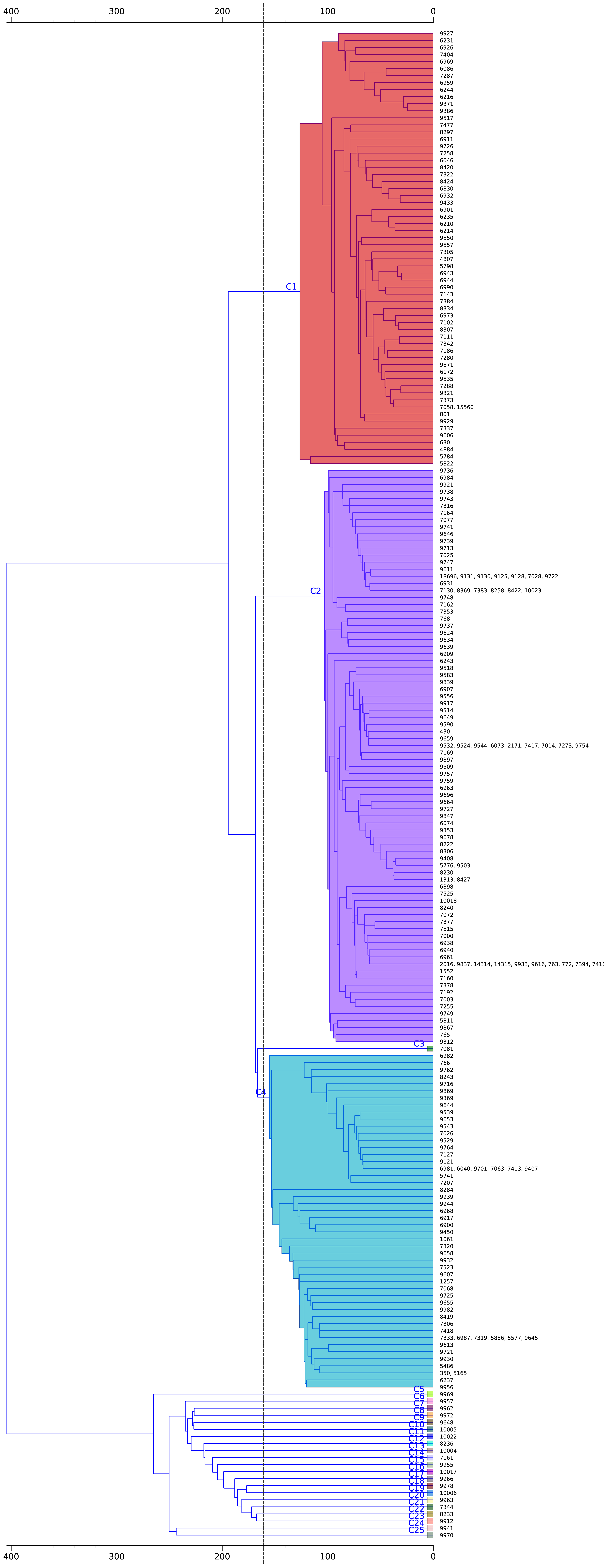

### Data S4

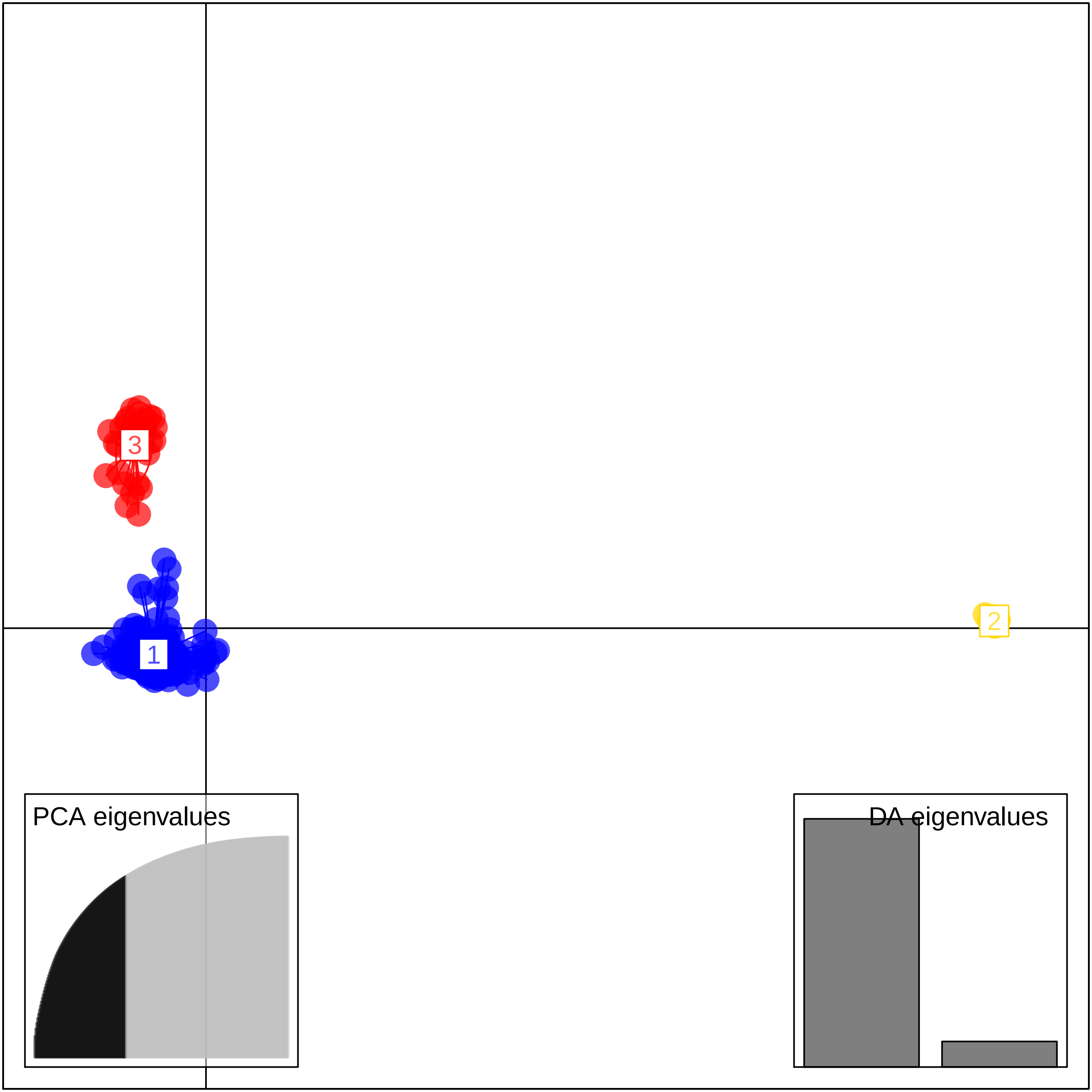

### Data S5

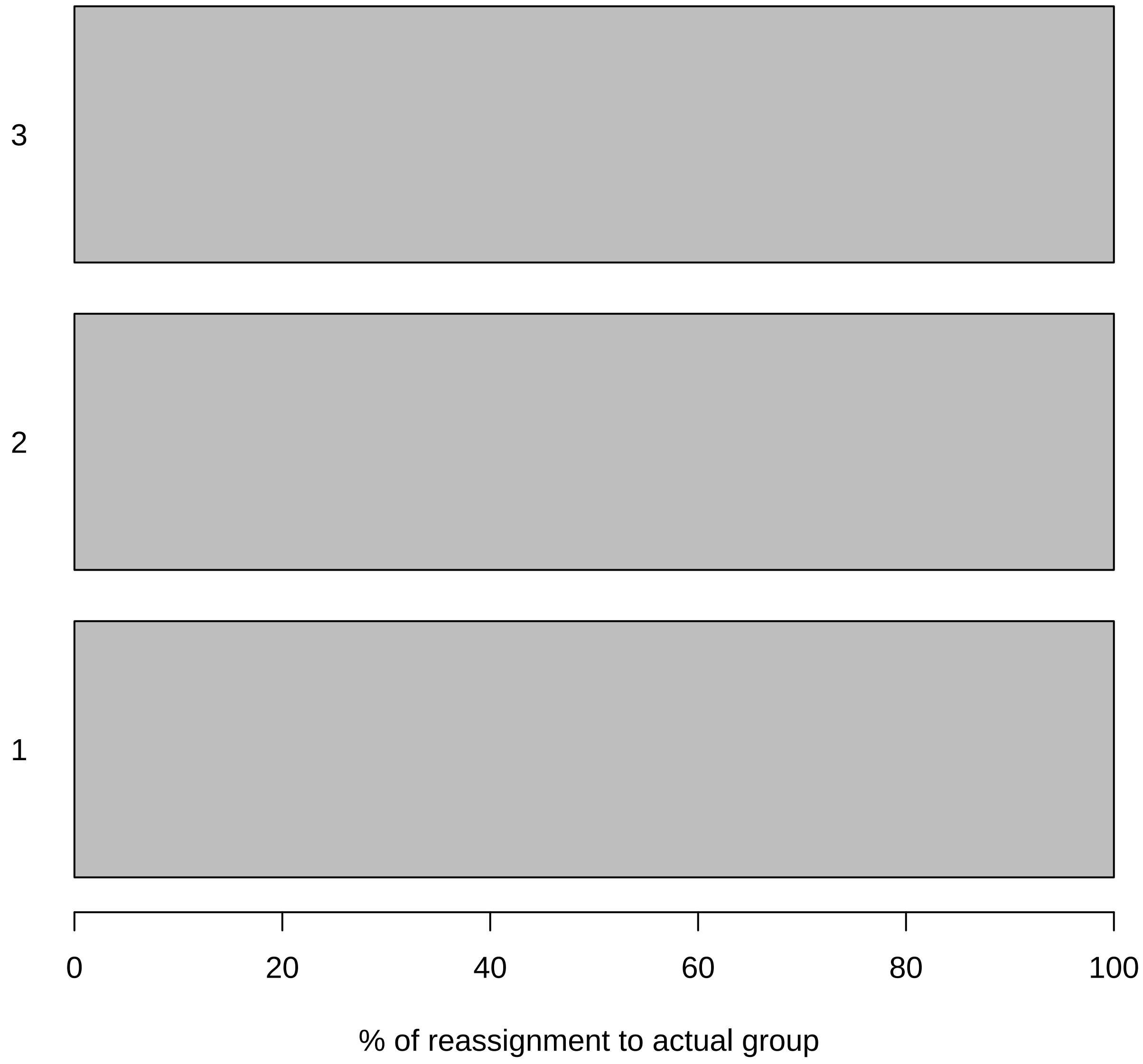

### Data S6

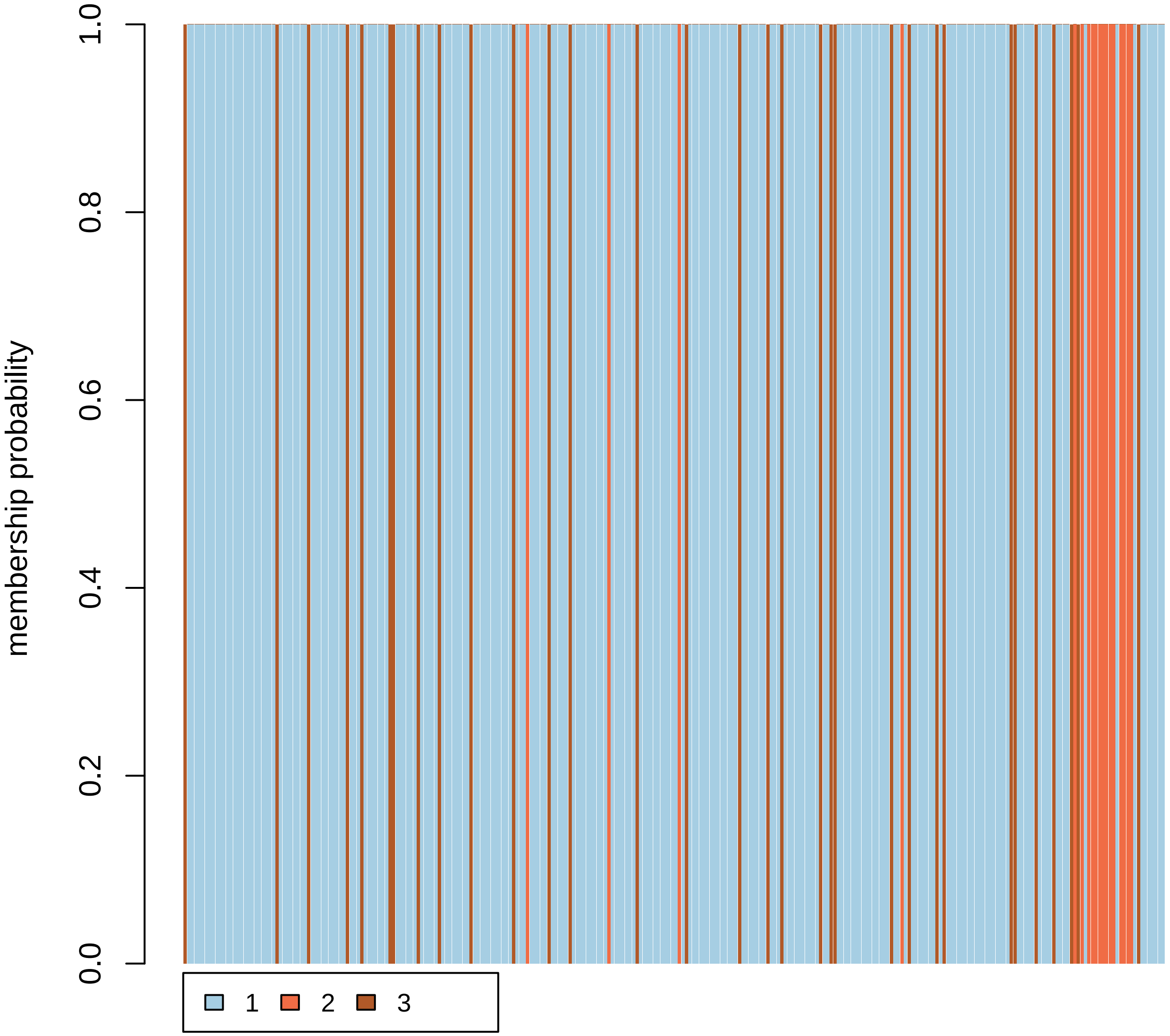
